# Physiological trade-offs drive the archaeal dominance and carbon turnover in deep subsurface

**DOI:** 10.64898/2026.05.21.726758

**Authors:** Jialin Hou, Lewen Liang, Liuyang Li, Weikang Sui, Liang Dong, Longhui Deng, Haining Hu, Zijun Wu, Lin Zhang, Orit Sivan, James A. Bradley, Fengping Wang

**Affiliations:** State Key Laboratory of Submarine Geoscience; Key Laboratory of Polar Ecosystem and Climate Change, Ministry of Education; Shanghai Key Laboratory of Polar Life and Environment Sciences; and School of Oceanography, Shanghai Jiao Tong University, Shanghai, China; State Key Laboratory of Microbial Metabolism, School of Life Sciences and Biotechnology, Shanghai Jiao Tong University, Shanghai, China; State Key Laboratory of Marine Geology, Tongji University, Shanghai, China; Department of Earth and Environmental Sciences, Ben-Gurion University of the Negev, Beer Sheva, Israel; Aix Marseille Université, Université de Toulon, CNRS, IRD, MIO, Marseille, France; School of Biological and Behavioural Sciences, Queen Mary University of London, London, UK

## Abstract

Marine sediments host a vast deep biosphere, yet how microorganisms persist under severe energy limitation and govern long-term organic carbon (OC) preservation remains poorly understood. Here we show that archaea, primarily *Bathyarchaeia*, systematically displace bacteria with depth and form net growth zones across East China Sea shelf deep sediments. Multi-omics analyses and bioenergetic modelling reveal that this transition is driven by sustained archaeal metabolism of diverse recalcitrant OC compounds, and a physiological trade-off that prioritizes cellular maintenance over growth, minimizing mortality in deep sediments. This strategy triggers a fundamental shift in sedimentary carbon turnover: from rapid bacterial degradation of labile OC near the surface to persistent archaea-driven turnover of recalcitrant OC at depth. We estimate that *Bathyarchaeia* mediate ∼77% of total OC degradation after 1,000 years of burial, corresponding to ∼18% (∼11.4 Pg C) of millennial OC degradation in global shelf sediments. These findings identify subsurface archaea as key microbial regulators of long-term OC preservation and reveal how physiological trade-offs sustain life and carbon turnover in the energy-limited deep biosphere.

## Main

Marine sediments represent one of Earth’s largest microbial habitats, harboring an estimated 2.9–5.4 × 10^29^ cells^1–3^. These subseafloor microbes drive biogeochemical transformations over geological timescales, ultimately governing long-term carbon sequestration and climate dynamics^4–6^. In the deep subsurface, however, microbial life operates under extreme energy limitation, with biomass turnover times extending from decades to millennia^7,8^. The systematic power-law decline of total cell abundance with burial depth has traditionally been interpreted as a passive winnowing process^1,2^, where only a small fraction of surface-derived microbes survive burial and persist in metabolic stasis or dormancy^9–11^. However, growing evidence of microbial growth at specific geochemical interfaces and physiological activities of *in vitro* cultivation challenges this monotonic decay view^12–14^, suggesting that certain lineages may sustain active growth in the deep subsurface and persist over geological timescales. Reconciling these two divergent views requires a shift in perspective: from treating subseafloor microbial communities as physiologically uniform to recognizing lineage-specific strategies that govern survival and persistence during burial. Archaea and Bacteria differ fundamentally in cellular architecture, physiology, and metabolism^15,16^, and have been hypothesized to follow distinct adaptation strategies and ecological succession trajectories in these energy-limited environments^17^.

Although estimates of archaeal abundance in marine sediments remain debated^3,12,18,19^, accumulating evidence indicates that subseafloor Archaea are widespread and ecologically significant, particularly on continental shelves, where they may comprise ∼37% of global sedimentary cells^20^. Among these, the class *Bathyarchaeia* is one of the most prominent uncultivated archaeal lineages, distinguished by its global distribution, high abundance, and metabolic versatility in carbon utilization^21–23^. Recently, a globally distributed *Bathyarchaeia* lineage was shown to exhibit mixoorganotrophic capabilities, simultaneously assimilating recalcitrant organic compounds (such as lignin) and CO_2_ as carbon sources^24–27^. These traits highlight the ecological significance of *Bathyarchaeia* in the deep biosphere, however, the specific metabolic and physiological mechanisms enabling their adaptation and persistence under the energy-limited conditions in the deeply buried sediments are not yet understood. This mechanistic gap limits our ability to quantitatively assess their contribution to global carbon cycling over geological timescales.

Here, we present high-resolution depth profiles of microbial abundance, diversity, metabolic activity and geochemistry from East China Sea (ECS) shelf sediments – a globally representative system characterized by high sedimentation rates and active organic carbon (OC) mineralization^28,29^. Our results reveal a systematic microbial succession from bacterial to archaeal dominance, driven primarily by the sustained net growth of *Bathyarchaeia* at depth. Multi-omic analyses indicate that this growth is fueled by active microbial remineralization of recalcitrant organic compounds. By integrating these observations into a millennial-scale bioenergetic model, we show that *Bathyarchaeia* adopt a distinct physiological trade-off, characterized by ultralow maximum growth rates and preferential energy allocation to cellular maintenance. This strategy reduces biomass loss, enabling net population persistence and sustained activity in the energy-limited deep subsurface. It drives a fundamental shift in the carbon cycling of marine sediments, from rapid, bacteria-dominated processing of labile OC near the surface, to persistent, archaea-mediated degradation of recalcitrant OC at depth. Our findings provide a mechanistic framework for understanding the ecological significance of Archaea in the deep biosphere, highlighting their pivotal, yet previously underappreciated role in regulating long-term organic carbon sequestration in marine sediments.

### Subseafloor microbial succession shifts from bacteria to archaeal dominance at depth

We collected six sediment cores (A2, A3, A5, B1, B2 and B3; length of 460-600 cm) from the Yangtze and Ou River estuaries (water depth 15-58 m) (Fig. 1a and Supplementary Table 1), where riverine inputs drive rapid sedimentation rates (mean rate of 3.42 cm/yr for A3^30^) and a high proportion of terrestrially-derived OC content (34-71% on average; Supplementary Fig.1). Four of these cores penetrated the sulfate-methane transition zone (SMTZ), while sulfate remained undepleted in two cores (Supplementary Fig.2). Quantitative PCR normalized with 16S rRNA gene copy number (archaea: 1.7; bacteria: 5.3^31^) revealed that archaea accounted for 48-65% (55% on average) of the total prokaryotic cells across the six ECS shelf cores (Fig.1b; Supplementary Table 2). This proportion substantially exceeds the ∼40% average reported for marginal sediments^20^, aligning with the observed negative correlation between archaeal abundance and water depth. Given that continental shelves (water depths < 150m) harbor at least one-third of microbial cells in global marine sediments^1^, this finding indicates that shelf archaea likely represent a dominant component of the global deep biosphere.

**Fig. 1.**
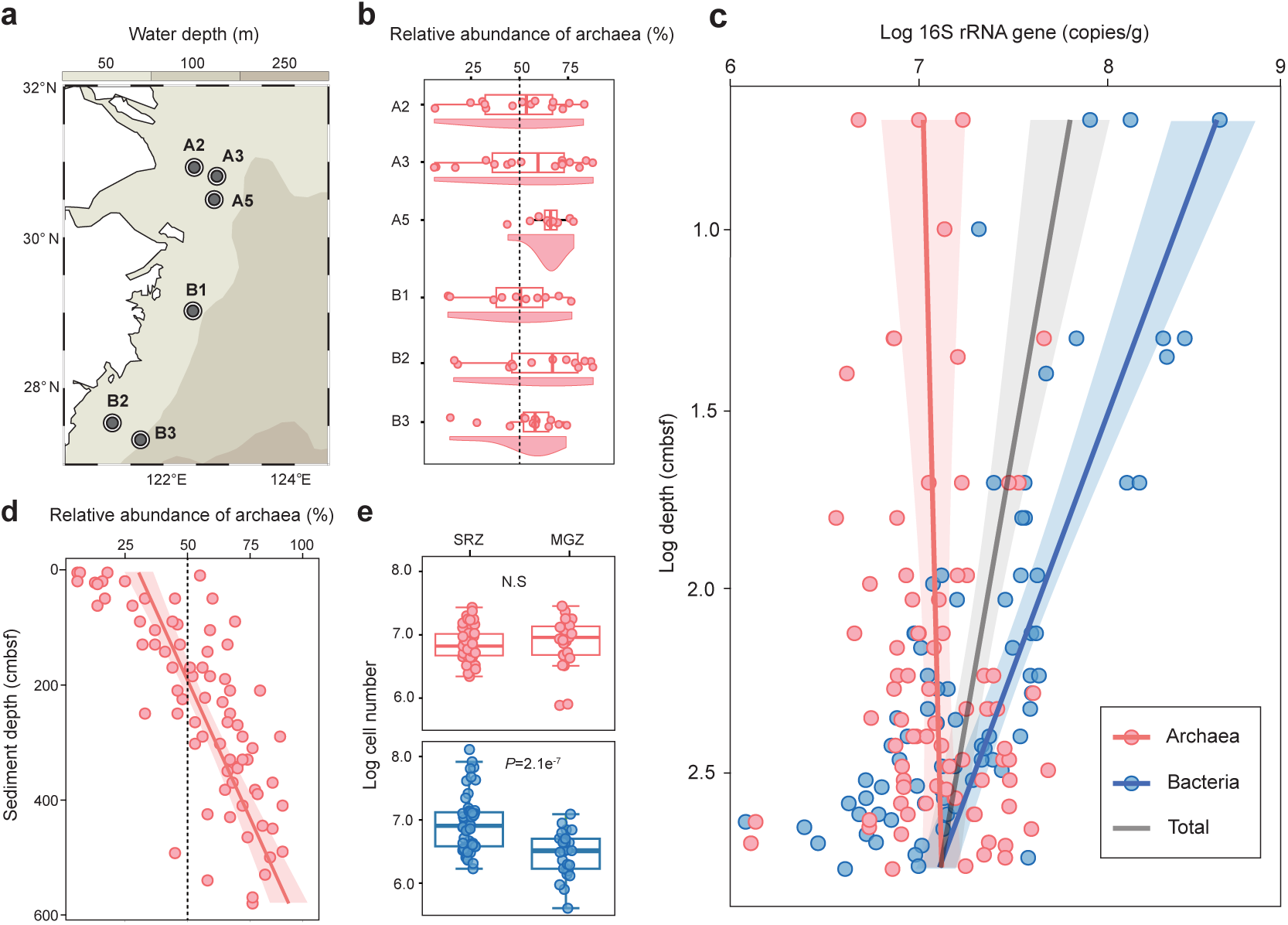
Microbial abundance and vertical distribution in the East China Sea (ECS) shelf sediments. **a,** Geographic locations of six gravity sediment cores collected from the ECS continental shelf. **b**, Raincloud plots showing the vertical distribution of archaeal relative abundance within each core. The dashed vertical line marks the 50% threshold. **c,** Vertical profiles of archaeal, bacterial and total prokaryotic 16S rRNA gene abundance across the six cores, with regression fits as a function of the sediment depth (cmbsf, centimeter below the seafloor): red, log (archaeal 16S rRNA gene abundance) = 6.99 + 0.05 log(depth), R^2^ = 0.01; blue, log (bacterial 16S rRNA gene abundance) = 9.0 – 0.71×log(depth), R^2^ = 0.57; grey line, log (total prokaryotic 16S rRNA genes) = 8.03 – 0.33×log (depth), R^2^ = 0.14. Shaded areas represent the 95% confidence intervals for the regression fits. **d,** Vertical distribution of archaeal cell relative abundance across the six cores, with the regression line: archaeal cells relative abundance (%) = 30.3 + 0.1034 × depth (cmbsf), R^2^=0.59. The overall average archaeal proportion exceeds 50% threshold (dash line) at depths deeper than 190 cmbsf. **e,** Comparison of estimated microbial cell numbers between the sulfate reduction zones (SRZ) and methanogenic zones (MGZ) across the six cores. Archaea maintain stable populations across geochemical transition zones (N.S., not significant), while bacterial populations exhibit a significant decline (P = 2.1×e^-7^). Note: In **d** and **e**, bacterial and archaeal cell numbers were calculated based on the average 16S rRNA gene copy numbers in archaeal and bacterial genomes (archaea: 1.7 copies, bacteria, 5.3 copies; rrnDB version 5.8)^31^,respectively. All regression statistics and geochemical zone definitions (Supplementary Fig.2) are based on the combined datasets from all samples of the six ECS cores.

**Fig. 2.**
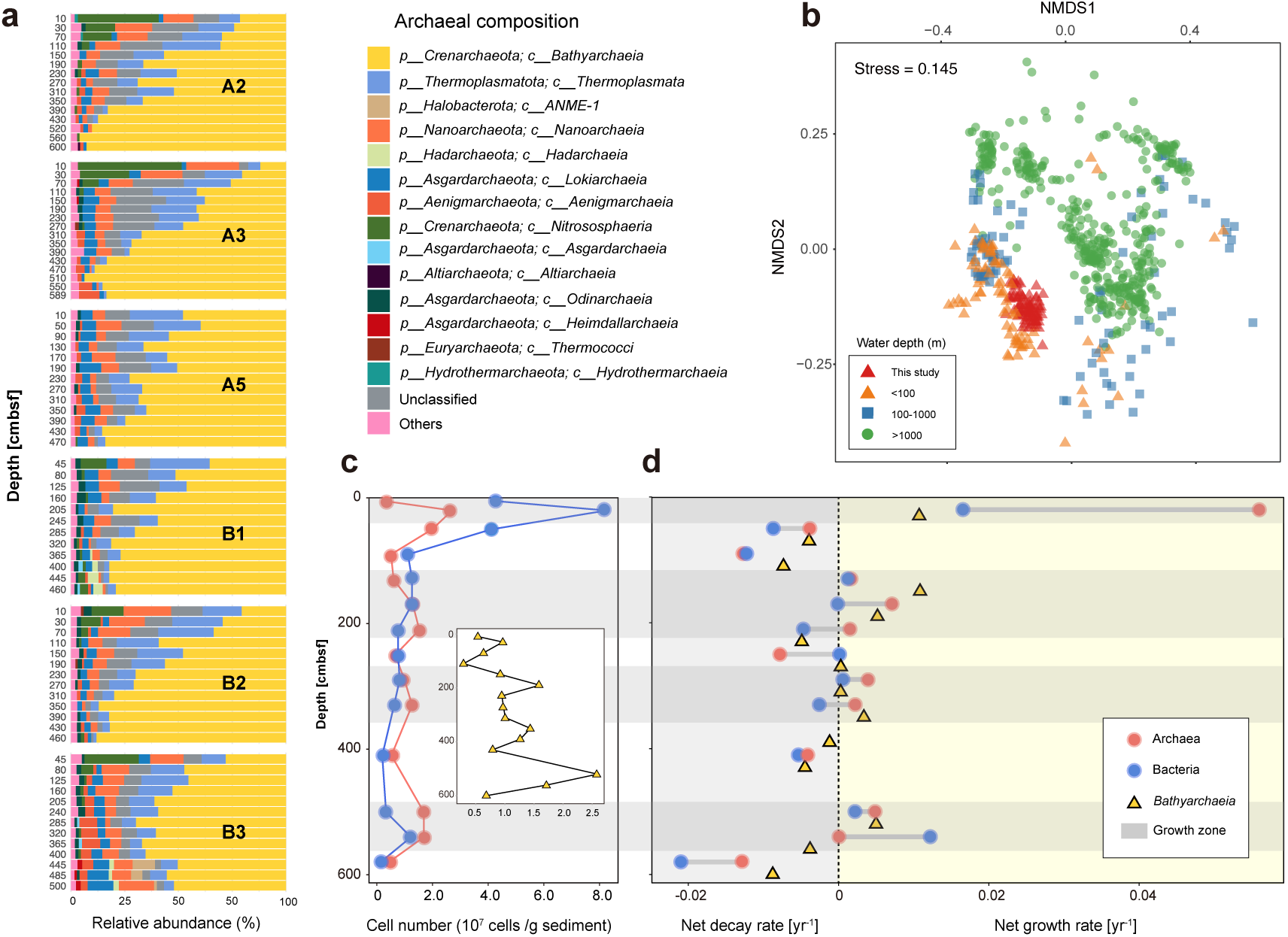
Microbial diversity and net growth rates in ECS sediment. **a**, Vertical distribution of archaeal community composition at the class level across the six sites (A2, A3, A5 B1, B2, and B3). The relative abundance of Class *Bathyarchaeia* (yellow bars) increases with depth, constituting the predominant archaeal lineage in deep sediments. Sediment depths represent the lower sample boundaries. Detailed depth specific to each sample is listed in the Supplementary Table 3. **b**, Non-metric multidimensional scaling (NMDS) analysis of 16S rRNA gene sequences comparing the ECS samples from this study (red triangles) with those from global marine sediment datasets. The NMDS plot is based on the Bray-Curtis dissimilarities of a rarefied ASV table (10,000 ASVs per sample). The global datasets (692 samples in total) were compiled from previous publications^32–35^ that used the same universal primer set 515F/806R^53^ (Supplementary Table 4). **c**, Depth profiles of bacterial (blue) and archaeal (red) cell abundances at core A2, quantified by 16S rRNA gene qPCR normalized using domain-specific average gene copy number per genome (as described in Fig.1). The subpanel shows the depth profile of *Bathyarchaeia* cell abundance (yellow triangle). **d**, Net growth and decay rates (yr^-1^) for Archaea, Bacteria and *Bathyarchaeia* throughout core A2, estimated using the fraction of read abundance multiplied by cell abundance (FRAxC)^13^. Positive values indicate net growth, while negative values represent net decay; grey-shaded bands highlight the specific subsurface microbial “growth zones”, where Archaea and *Bathyarchaeia* exhibit sustained net growth.

Total prokaryotic 16S rRNA gene abundance declined logarithmically with sediment depth in the ECS shelf cores (Fig. 1c), consistent with established global subseafloor cell distributions^1,20^. However, we observed divergent trajectories in the subsurface microbial communities: bacterial abundance decreased with sediment depth, whereas archaeal abundance increased in deeper layers (Fig.1c). Consequently, archaea dominated (>50%) in sediments deeper than ∼190 cm below the seafloor (cmbsf) across all ECS cores (Fig. 1d). This divergence reflects contrasting population dynamics among bacteria and archaea across geochemical zones: bacterial abundance declined significantly from the sulfate reduction zone (SRZ) to the methanogenic zone (MGZ), whereas archaeal populations remained relatively stable (Fig.1e). These findings suggest that Archaea likely employ distinct physiological strategies enabling their adaptation and dominance in the deep subsurface. Together, our results demonstrate a progressive subseafloor archaeal succession that is decoupled from the typical power-law decline of microbial cells in marine sediments, thus highlighting the key ecological roles of Archaea in the deep biosphere.

### *Bathyarchaeia* growth drives the deep archaeal dominance

Archaeal communities across the six ECS cores showed highly similar composition, predominantly composed of the phylum *Crenarchaeota*, followed by *Asgardarchaeota*, *Nanoarchaeota* and *Aenigmarchaeota* (Fig.2a; Supplementary Table 3), consistent with global surveys of anoxic marine sediments^32^. Specifically, the class *Bathyarchaeia* comprised >90% of the phylum *Crenarchaeota* across all samples, with an increasing proportion at greater sediment depths, and reaching ∼95% of the total archaeal community in core A2 at 560 cmbsf (Fig.2a), underscoring the predominance of *Bathyarchaeia* in deep shelf sediments. We compared our results with community profile data from globally distributed sediment samples that used identical primers^32–35^ (692 samples in total, Supplementary Table S4). Beta-diversity analysis showed that ECS microbial communities clustered with other continental shelf sediments (water depth < 100 m) globally, yet diverged from their deep-sea counterparts (water depth > 1,000 m) (Fig. 2b). This biogeographic pattern indicates that the ECS cores represent typical continental shelf sediment communities, where archaeal predominance, particularly *Bathyarchaeia*, constitutes a hallmark feature of microbial diversity in global continental shelf sediments. The ubiquity and dominance of *Bathyarchaeia* has been documented previously^21,23^, however previous studies primarily focused on estuary, mangrove and other shallow sediments^36–38^. Here, we show an increasing predominance of *Bathyarchaeia* in deeper shelf sediments, suggesting that this archaeal lineage possess specialized adaptive strategies enabling their dominance in the global deep subsurface.

To determine whether the observed increase in archaeal abundance with burial depth reflects active archaeal growth or differential mortality relative to bacteria (Fig. 2c), we quantified the net growth rates of archaeal, bacterial, and *Bathyarchaeia* populations in core A2. By integrating qPCR with 16S rRNA gene profiling through FRAxC (the fraction of read abundance multiplied by cell abundance) analysis^13^ (Supplementary Table 5), we identified peak net growth rates for bacteria and archaea within the upper 30 cmbsf, corresponding to their maximum cell densities (Fig. 2d). This near-surface growth zone aligns well with previous observations from estuarine sediments^13^, indicating that the sediment-water interface serves as a critical niche and selective filter for subseafloor microbial populations. We infer that the capability for persistence and proliferation in deep sediments is established during initial deposition, rather than acquired through *de novo* adaptive evolution during long-term burial^9^.

Although overall cell numbers and growth rates substantially diminished below 100 cmbsf, we detected multiple net microbial growth zones at depth, extending to ∼500 cmbsf (Fig.2d; Supplementary Table 5), indicating that active cell proliferation persists in deep shelf sediments. This finding challenges the prevailing paradigm of a deep biosphere dominated by microbial dormancy and ultra-slow biomass turnover^7,8,11^. Notably, archaeal growth rates consistently exceeded those of bacteria in deep sediment growth zones, particularly for *Bathyarchaeia* (Fig.2d), consistent with the divergent vertical distribution patterns of Archaea and Bacteria (Fig.1c). Although *Bathyarchaeia* are known to be metabolically active in marine sediments^39,40^, our results demonstrate their *in situ* population growth at depth, and explain the increased archaeal dominance in deeply buried shelf sediments.

### Active recalcitrant OC utilization of *Bathyarchaeia* at depth

To investigate the metabolic mechanisms underlying the archaeal predominance in deep shelf sediments, we performed integrated metagenomic and metatranscriptomic analyses across five depths (10, 150, 190, 390 and 600 cmbsf) in the core A2, which shows pronounced archaeal dominance in deep layers (Fig. 2a). Our results show that subseafloor microbes encoded and actively transcribed key genes for diverse carbon metabolic pathways across all depths, including the Wood-Ljungdahl pathway, anaerobic fermentation, aromatic degradation, C1 compounds and acetate metabolism, enzymes for protein and carbohydrate decomposition (Fig. 3a; Supplementary Table 6). These profiles indicate the sustained physiological and carbon metabolic activities of microbial communities in shelf sediments long after their deposition on the seafloor, consistent with documented metabolic activities in deep Peru Margin sediments^41–43^. Notably, despite the general decline in microbial potential and transcriptional activity with burial depth, these shelf communities exhibit elevated expression levels for most of these key genes in the deeper SMTZ (390 cmbsf) and MGZ (600 cmbsf) compared to shallower sediments (from 10-190 cmbsf) (Fig. 3a). This finding indicates that surviving lineages are physiologically adapted to energy limitation through enhanced transcriptional efficiency, thus supporting their persistent contribution to the carbon cycle in the deep subsurface.

**Fig. 3.**
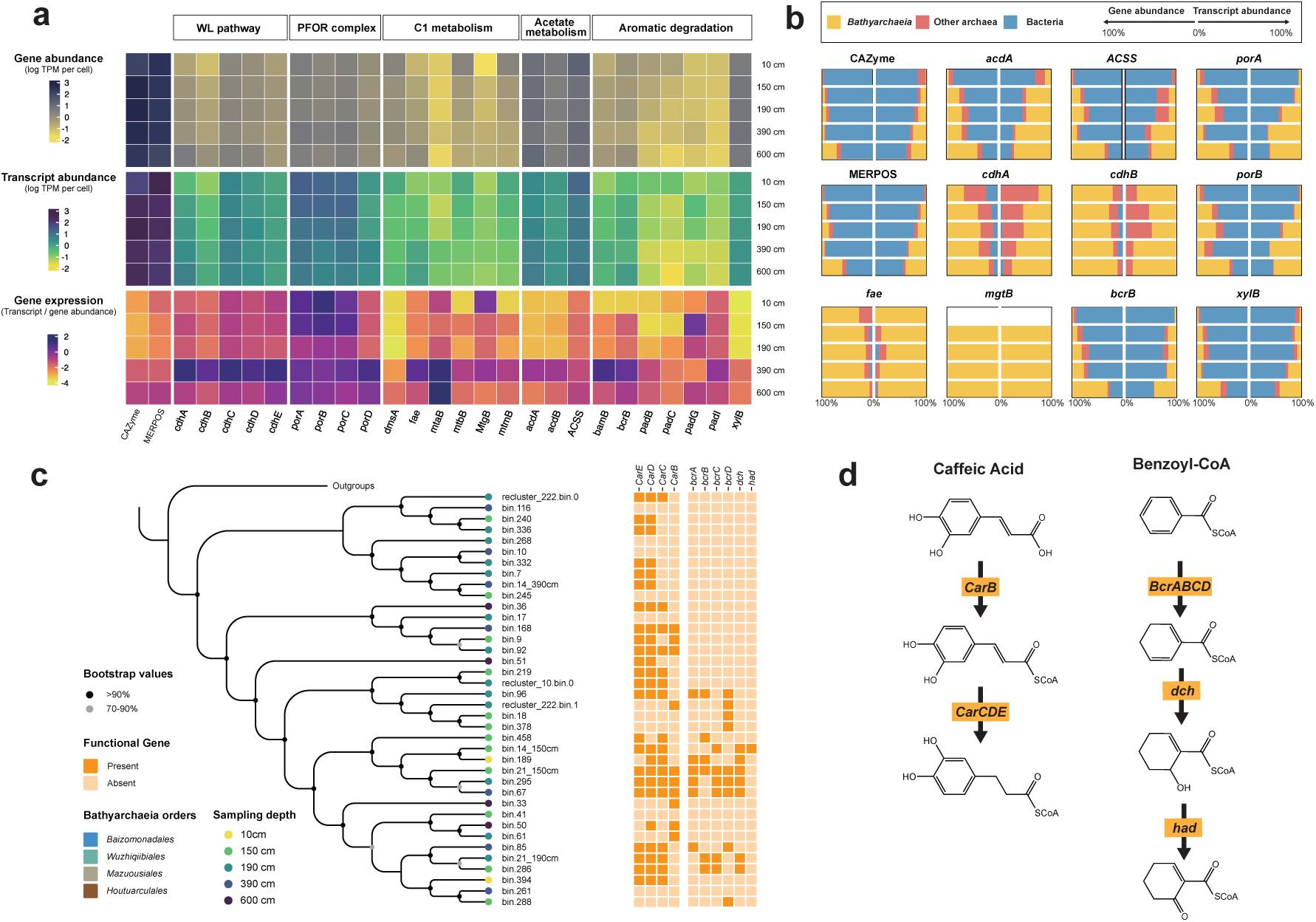
Carbon metabolic gene profiling of *Bathyarchaeia* in site A2. **a**, Depth profiles of gene abundance, transcript abundance, and expression level (ratio of transcript to gene abundance) for marker genes in carbon metabolic pathways, including Wood-Ljungdal (WL) pathway, pyruvate ferredoxin oxidoreductase (PFOR) complex, C1 compounds metabolism, acetate metabolism and aromatic degradation (Supplementary Table 6). **b,** Taxonomic composition of *Bathyarchaeia*, other archaea, and bacteria contributing to key marker genes across five sediment depths, visualized as mirrored barplots for gene (left) and transcript (right) abundances. **c,** Phylogenetic placement of B*athyarchaeia* MAGs reconstructed from five depths in core A2, along with the distribution of key genes for aromatic compound degradation among different *Bathyarchaeia* lineages. **d**, The key genes for anoxic caffeic acid and benzoate-CoA degradation pathways identified in the *Bathyarchaeia* MAGs. Abbreviations: *carB*, caffeyl-CoA synthetase, *carC*, caffeyl-CoA reductase; *carDE*, electron transfer flavoprotein A and B; *bcrABCD*, benzoyl-CoA reductase subunit ABCD; *dch*, cyclohexa-1,5-dienecarbonyl-CoA hydratase; *had*, 6-hydroxycyclohex-1-ene-1-carbonyl-CoA dehydrogenase; MEROPS, the Peptidase Database; CAZyme, Carbohydrate-Active enZymes Database. *Mto_m*, gene encoding the novel methyltransferase found in *Methermicoccus shengliensis*; *Mto_a*, gene encoding the novel methyltransferase found in *Archaeoglobus fulgidus*; *MtgB*, gene encoding the novel methyltransferase found in *Bathyarchaeia* lineages^27^.

As the dominant archaeal lineage in shelf sediments, *Bathyarchaeia* play a central role in driving the turnover of diverse organic compounds, including carbohydrates, proteins, xylose, methylated, and aromatic compounds, likely through fermentation and acetate metabolism (Fig.3b). This metabolic versatility is consistent with their known capabilities to utilize diverse biopolymers^44^, proteins^45^, and lignin as carbon and energy sources^24,26^. For most key genes, *Bathyarchaeia* dominated both the gene abundance and transcriptional activity within Archaea, with their relative contribution increasing with sediment depth (Fig.3b). Notably, *Bathyarchaeia* exclusively encode *mtgB*, the key gene of a novel methyltransferase system involved in transferring methoxyl groups from lignin^27^. We detected this gene across all four anoxic depths (150-600 cmbsf), with transcripts specifically assigned to *Bathyarchaeia* (Fig.3b). Moreover, this gene showed markedly higher expression relative to other methyltransferase-encoding genes in the deep SMTZ and MGZ (Supplementary Fig.3). These findings indicate that *Bathyarchaeia* actively metabolize diverse organic compounds in deep shelf sediments, with their ecological importance increases at deeper depth. These metabolic traits not only sustain their predominance in the global deep biosphere, but also reveal a critical, yet previously underappreciated roles of archaea in the deep subsurface carbon cycle.

To elucidate the carbon metabolic mechanisms of *Bathyarchaeia*, we recovered 38 high- and medium-quality *Bathyarchaeia* metagenome-assembled genomes (MAGs) from five depths in core A2, which are taxonomically affiliated within the orders *Baizomonadales*, *Wuzhiqiibiales*, *Houtuarculales* and *Mazuousiales* (Fig. 3c). Most of these MAGs encoded key genes involved in the anaerobic degradation of aromatic compounds and phenolic acids, including *bcrABCD* (benzoyl-CoA reductase subunit ABCD), *dch* (cyclohexa-1,5-dienecarbonyl-CoA hydratase), *had* (6-hydroxycyclohex-1-ene-1-carbonyl-CoA dehydrogenase), *carBCDE* (caffeyl-CoA synthetase/reductase and electron transfer flavoprotein) (Fig 3c,d). These findings indicate that these *Bathyarchaeia* lineages are capable of utilizing recalcitrant organic compounds throughout the ECS sediment core. Specifically, ATP-dependent benzoyl-CoA reductase (*bcrABCD*) catalyzes the anaerobic cleavage of aromatic rings, a central step in anaerobic aromatic degradation^46^. Although *bcrABCD* genes have been identified in several *Bathyarchaeia* MAGs from estuarine sediments^37^, our results further demonstrate that these genes are exclusively encoded by *Baizomonadales*, which dominates both core A2 and global anoxic marine sediments^22,24^. Furthermore, GC-MS analysis confirmed the presence of diverse aromatic compounds throughout the core A2, with concentration profiles generally declining with depth (Supplementary Fig.4). These findings indicate that active anaerobic aromatic degradation is a common metabolic strategy for the *Bathyarchaeia* lineages to exploit recalcitrant organic substrates in deep shelf sediments, supporting their ecological advantages and population growth in the energy-limited deep subsurface.

**Fig. 4.**
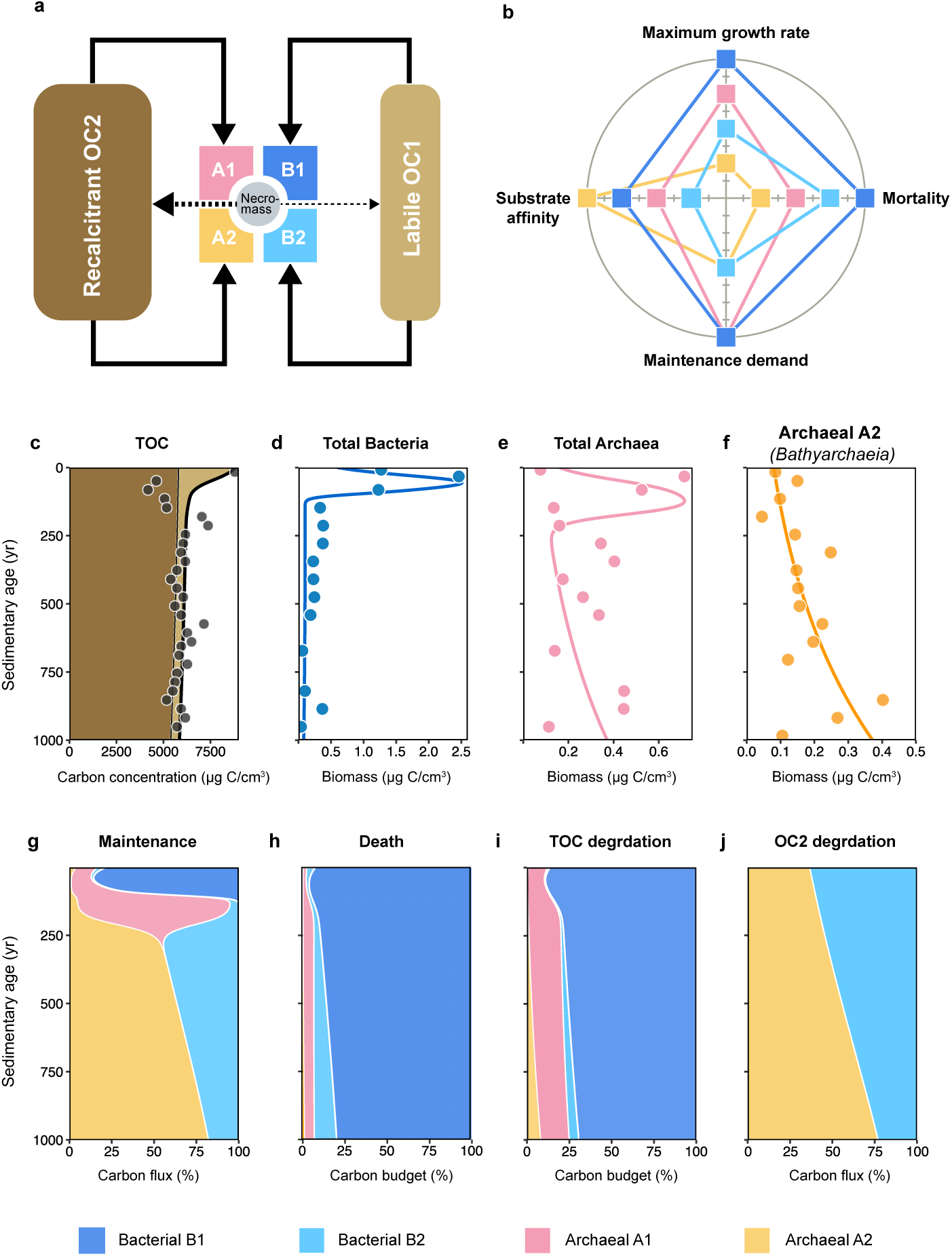
Integrated bioenergetic model of microbial succession and OC degradation in the ECS shelf sediments. **a,** Schematic diagram of the bioenergetic model, which explicitly couples the degradation of two OC pools (labile OC1 and recalcitrant OC2) with four microbial guilds (B1, B2, A1 and A2) defined by distinct substrate preferences and life-history strategies. Necromass produced from dead microbial cells is partially recycled into the OC pool, mostly as labile OC1. **b**, The divergent physiological trade-offs among the four microbial guilds. The radar chart illustrates their distinct life-history strategies across four parameters: maximum growth rate, mortality, maintenance demand and substrate affinity. Line colors correspond to the different microbial guilds in Fig.4a. Note that the substrate affinity of B1 and A1 are parameterized to labile OC1, whereas B2 and A2 specifically target to recalcitrant OC2. The values in the radar chart are presented on a relative scale to compare the physiological trade-off among microbial guilds, specific parameters used in the model are detailed in the Supplementary Methods. Observed (dots) and simulated (lines) total OC concentration profile along burial age (**c**) in which dark and light green represent the labile and recalcitrant OC pools, respectively. **d-f,** Depth profiles of observed (dots) and simulated (lines) microbial biomass of bacteria (**d**), archaea (**e**) and Bathyarchaeia (archaeal guild A2; **f**). **g-h,** Simulated carbon fluxes and budgets among the four microbial guilds. Stacked colored areas represent the relative contributions of the four guilds to the total carbon fluxes allocated to maintenance demand (**g**), necromass produced from dead cells (**h**). **i,** Relative contributions of four microbial guilds to the total OC degradation budget. **j,** degrading rate of recalcitrant OC driven by bacterial guild B2 and Bathyarchaeia (archaeal guild A2).

### Physiological trade-offs dictate *Bathyarchaeia* growth and the deep carbon cycle

To resolve mechanistically how *Bathyarchaeia* achieve net growth in the deep subsurface, we developed a constrained bioenergetic model that explicitly couples OC degradation with microbial population dynamics during millennial-scale burial. The model incorporates two OC pools (labile OC1 and recalcitrant OC2) and four microbial guilds (Bacteria: B1 and B2; Archaea: A1and A2) with distinct substrate preferences and life-history strategies (Fig.4a). Specifically, bacterial guild 𝐵_1_represents copiotrophs exclusively targeting labile OC1 for rapid growth, at the cost of high maintenance demands and mortality rates. The archaeal guild A1 adopts a similar but more moderate strategy to balance growth and maintenance, resulting in its persistence to greater depths than B1. By contrast, the bacterial guild B2 and archaeal guild A2 compete for recalcitrant OC2 (Fig.4b). Calibrated to *in situ* rates of recalcitrant OC degradation (Fig.3a), guild A2 serves as a proxy for *Bathyarchaeia*. It exhibits the lowest maximum growth rate, growth yield, and mortality among all guilds, with a higher substrate affinity (lower half-saturation constant) than B2. Constraining the model with OC and biomass profiles from core A2 (Fig.4c-j), we tested whether microbial growth in the deep subsurface is governed by vertical geochemical gradients or distinct physiological trade-offs arising during long-term burial.

Our simulations revealed a rapid OC mineralization near the sediment surface, where ∼77% of labile OC is consumed within the first ∼100 years of burial, accounting for ∼26% of total OC loss over the millennial timescale (Fig.4c). This near-surface mineralization hotspot is primarily governed by the activity of the bacterial guild B1, which has the highest maximum growth rate and substrate affinity among all four guilds (Fig.4b), thus enabling their rapid population increase and intensive OC degradation shortly after deposition on the seafloor (Fig.4d; Extended Data Fig.1). However, this copiotrophic strategy is intrinsically coupled to higher maintenance demands and mortality (Fig.4b and g). As a consequence, the bacterial guild B1 and the bacteria community as a whole, is highly sensitive to substrate depletion, thus undergoing a rapid collapse in population once the labile OC concentrations fall below the threshold required to sustain their high maintenance demands (Fig.4d; Extended Data Fig.1). By contrast, the archaeal guild A1 avoids direct competition with the B1 guild in shallow sediments by adopting a lower substrate affinity, slower growth rate, and reduced maintenance requirements (Fig.4b), enabling persistence at greater depths and reaching maximum population after ∼120 years of burial (Fig.4e; Extended Data Fig.1). Overall, surface sediments thus function as a copiotrophic bacteria-driven hotspot of labile OC turnover, playing a pivotal role in early-stage carbon preservation in shallow sediments. By selectively removing the more labile components, microbial activity in surface sediments reshapes subsurface OC reactivity, effectively funneling a disproportionately recalcitrant OC pool towards deep preservation and long-term cycling over geological timescales.

In contrast to the rapid, bacteria-driven turnover of labile OC near the sediment surface, our simulations reveal that recalcitrant OC degradation proceeds at substantially lower rates but increases persistently with depth throughout the modeled timescale (Fig.4c; Extended Data Fig.2). This process of long-term carbon remineralization is dominated by the archaeal guild A2, here defined as *Bathyarchaeia* (Fig.4f), whose contribution to the recalcitrant OC degradation rises from ∼38% to ∼59% with increasing burial depth (Extended Data Fig.2). In 1,000-year-old sediments, *Bathyarchaeia* account for ∼77% of the total OC degradation, cumulatively representing ∼8.2% of total OC mineralization over the millennial timescale (Extended Data Fig.3). By scaling these dynamics using an established global framework^5^, we estimate that *Bathyarchaeia* degrade ∼4.8 Pg C in global shelf sediments over 1000 years of burial, reaching a maximum mineralization rate of ∼39.2 Tg C/yr at millennial-scale depth (Extended Data Fig.4). Although previous incubation and genomic studies have demonstrated the capability of *Bathyarchaeia* to degrade recalcitrant compounds^26,27^, our model quantitatively illustrates their pivotal role as the primary microbial driver of recalcitrant OC turnover in shelf sediments, highlighting their substantial impact on the deep subsurface carbon cycling over geological timescales. Integrating these findings with the transient bacteria-driven OC mineralization in surficial sediments, these findings support a vertically stratified carbon cycling framework (Fig.5) where copiotrophic bacteria drive initial carbon degradation in the uppermost shelf sediments, and archaea, particularly *Bathyarchaeia*, regulate the long-term fate and reactivity of OC in deep marine sediments.

**Fig. 5.**
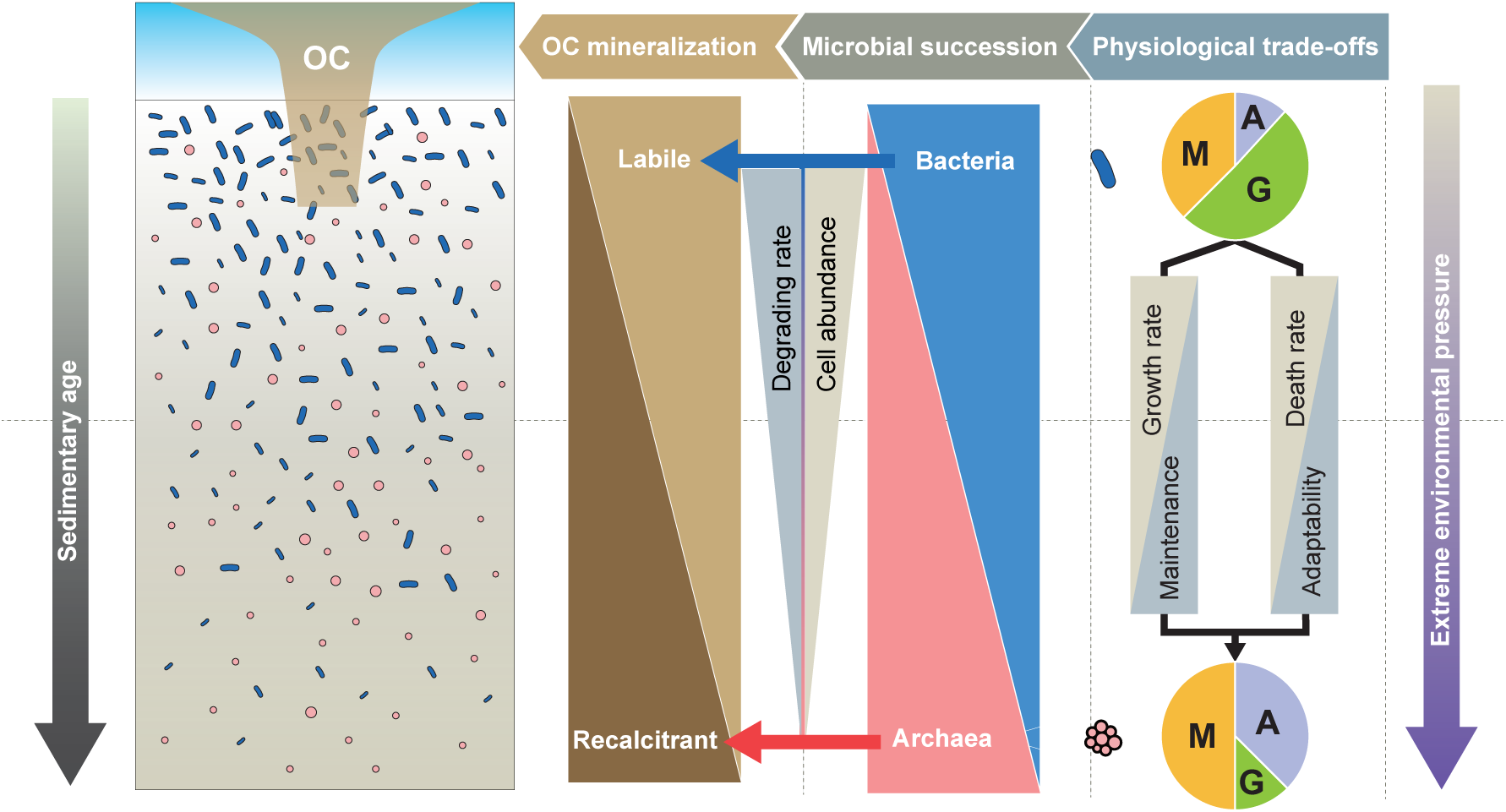
Conceptual framework of vertically stratified carbon cycling and microbial succession driven by physiological trade-offs in shelf sediments. Organic carbon (OC) reactivity and microbial community structure undergo profound transitions with sediment depth. In surface sediments, Bacteria (blue) drive a transient mineralization hotspot by rapidly consuming labile OC, whereas Archaea (pink) dictate the long-term fate and reactivity of the buried recalcitrant OC pool over geological timescales. These transition of OC degradation and microbial succession are underpinned by fundamental physiological trade-offs: whereas Bacteria prioritize rapid growth (G) with high energy investment near the surface, Archaea, primarily *Bathyarchaeia*, allocate more energy toward maintenance (M) and adaptation (A) by minimizing their growth rates in the deep subsurface.

The classic ecological theory defines oligotrophy through minimized maintenance energy requirements and slow growth to ensure persistence in resource-limited environments^7,8,47^. Our findings, however, suggest that *Bathyarchaeia* in deep shelf sediments are not typical oligotrophs. Rather, their ultra-slow growth rate provides a strategic bioenergetic advantage that allows for the preferential allocation of limited energy toward essential maintenance and adaptation in the extreme deep subsurface environment (Fig.5). This is also supported by their higher transcriptional activity of antioxidation systems in deeper sediments (Supplementary Fig.5), a key mechanism for microbial adaptation to multiple extreme conditions^48^. Consequently, this physiological trade-off effectively minimizes the mortality of *Bathyarchaeia*, as reflected by their extremely low death rate in the deep subsurface (Fig.4b) and minor contribution (<1%) to the necromass pool during millennial-scale burial (Fig.4h; Extended Data Fig.5). This strategy resembles the growth-survival trade-off of clinical pathogens^49^, which exhibit slow growth phenotypes inside hosts^50^. Previous studies suggest that microbial communities persist in the deep subsurface either in metabolic stasis or dormancy^8,11^, with succession primarily driven by selective survival of pre-adapted lineages, rather than *de novo* adaptation during burial^9,51^. However, the net growth of *Bathyarchaeia* observed in deep shelf sediments demonstrates that certain subsurface lineages, particularly Archaea, employ unique metabolic and physiological strategies to replicate and function in deep energy-limited environments. Our work indicates that microbial succession in the deep biosphere is not merely a passive winnowing process, but rather a dynamic physiological trade-off for long-term persistence. Together, this study reveals the physiological mechanisms underlying the ecological success of *Bathyarchaeia* and elucidates how these archaea survive in deep subsurface sediments and regulate the subsurface OC preservation over geological timescales.

## Materials and Methods

### Sample collection and geochemical analyses

Six gravity sediment cores (A2, A3, A5, B1, B2, B3) were collected from the East China Sea (ECS) continental shelf (Fig. 1a) during the cruise organized by International Center for Deep Life Investigation (IC-DLI) from 8 to 15 July, 2017. These cores were retrieved during the cruise at water depths of 15-58 m, with core length ranged from 460 cm to 600 cm (Supplementary Table 1). The sampling region is located offshore of eastern China and is characterized by mud deposits primarily derived from Yangtze River inputs^29^. Upon retrieval, porewater of sediment cores was extracted using Rhizon samplers (0.2-μm pore size; Rhizosphere). Pore water geochemical parameters (SO_42_^-^, Cl^-^, Ca^2+^, Mg^2+^, dissolved inorganic carbon (DIC) and δ^13^C-DIC), methane as well as total organic carbon (TOC) and δ^13^C-TOC were measured as described in the Supplementary Methods. Sediments for microbiological analysis were stored at -80°C until DNA and RNA extraction.

### DNA extraction, qPCR, 16S rRNA gene sequencing and analysis

DNA was extracted using the PowerSoil DNA Kit (MoBio Laboratories, CA, USA) following the manufacturer’s protocol and stored at -80 °C until further processing. Quantitative PCR (qPCR) assays for both bacterial and archaeal 16S rRNA genes were performed as previously described^52^, using bac341f (5′-CCTACGGGWGGCWGCA-3′) and 519r (5′-TTACCGCGGCKGCTG-3′) primer set for bacteria, Uni519f (5′-CAGCMGCCGCGGTAA-3′) and Arc908R (5′-CCCGCCAATTCCTTTAAGTT-3′) primer set for archaea. 20-μl reaction volumes containing 10 μl of 2X TB Green Premix Ex Taq II (Tli RNaseH Plus) (TaKaRa, Dalian, China), 0.4 μl of 50X ROX Reference Dye (TaKaRa), 0.8 μl of 10 μM primers (Sangon Biotech, Shanghai, China), and 1 μl of template DNA, with the final volume adjusted using autoclaved double-distilled water. Thermal cycling conditions consisted of an initial denaturation at 95 °C for 15 min, followed by 40 cycles of 95 °C for 30 s, an annealing step (56 °C for bacteria and 59 °C for archaea) for 30 s, and an extension step at 72 °C (30 s for bacteria and 45 s for archaea). All reactions exhibited an R^2^> 0.99 and amplification efficiencies of 90–110%.

The V4 region of 16S rRNA gene was amplified using barcoded 515F (5′-GTGYCAGCMGCCGCGGTAA-3′) and 806R (5′-GGACTACNVGGGTWTCTAAT-3′) universal primers^53^. Amplicons were sequenced on the Illumina NovaSeq 6000 platform (Illumina, CA, USA). Downstream sequence analyses were performed using QIIME2 (Version: 2022.2)^54^. Briefly, adapters were removed using Cutadapt v4.0^55^, and sequences were denoised, and clustered into amplicon sequence variants (ASVs) using DADA2^56^. Taxonomic assignments were performed using the classify-sklearn method in the q2-feature-classifier plugin^57^ against the SILVA 138 reference database^58^. Bacterial and archaeal ASVs were rarefied to 24,815 and 1,633 sequences per sample, respectively. We also collected published sequence data from global marine sediments from the literatures^32–35^, which were amplified using the same primer set (515F/806R) (Supplementary Table S4). All 692 samples were processed using the same bioinformatic pipeline as described above. To evaluate the distribution patterns of microbial communities in marine sediments, the ASV table was rarefied to 10,000 ASVs per sample, subsequent NMDS analysis was performed using the metaMDS function with Bray–Curtis dissimilarities based on the “Vegan” package^59^ in R.

### Metagenomic sequencing and analysis

Samples from five depths (10, 150, 190, 390, and 600 cm) of core A2 were selected for DNA extraction and metagenomic sequencing. Paired-end reads were generated on a Novaseq 6000 platform (Illumina, CA, USA). Subsequent reads processing and analyses were performed following a modified in-house pipeline from our laboratory. Briefly, raw reads were trimmed using Trimmomatic v0.39^60^, the resulting clean reads were assembled into contigs by MEGAHIT v1.2.9^61^ with the parameters -min-count 2 -k-min 41 -kmin-1pass -k-max 147 -k-step 10. Open reading frames (ORFs) were predicted from the contigs and translated into amino acid sequences using prodigal v2.6.3^62^ in -meta mode.

Functional genes were annotated using the DRAM v1.4.0^63^ against KEGG^64^, UniRef90^65^, PFAM^66^ databases with default parameters, followed by checked manually for specific key genes (Supplementary Table 6). The carbohydrate-active enzymes (CAZymes)^67^were further annotated using HMMER 3.3.2^68^ (E-value < 10−15) on CAZymes V10 database. Proteolytic enzymes and peptidases were annotated using Diamond blastp (v0.9.14) (E-value < 10^−15^ and coverage > 30%) against MEROPS database^69^. For each functional gene group (e.g. KO), clean reads were mapped to each gene using Bowtie2 v2.4.4^70^ under the -very-sensitive mode, which was further normalized to gene abundance per cell in each sample by dividing the median abundance of 10 single-copy conserved marker gene (K06942, K01889, K01887, K01875, K01883, K01869, K01873, K01409,K03106, and K03110) according the method described in Salazar’s work^71^.

Contigs longer than 1.5 kb, 3kb, and 5Kb were assigned for recovering metagenome assembled genomes (MAGs) using MetaBAT2 v2.2.15^72^ and Maxbin v2.2.7^73^ with default parameters. These MAGs were refined using RefineM v0.0.22^74^ and subsequently integrated into optimized, non-redundant bins using DAS Tool v1.1.4^75^. The quality of MAGs was estimated using CheckM v1.0.9^76^ with lineage-specific markers genes parameter, and assigned taxonomy by GTDB-tk^77^ based on the GTDB taxonomy (r220). The relative abundance of MAG was calculated using coverM v0.7.0^78^.

### RNA extraction, sequencing and metatranscriptomic analysis

In parallel, RNA from the same five depths of core A2 was extracted using the RNeasy PowerSoil Total RNA Kit (Qiagen, Germany) according to the manufacturer’s instructions. Metatranscriptomic libraries were generated using the NEB Next UltraTM Nondirectional RNA Library Prep Kit for Illumina (New England Biolabs, MA, USA) following the manufacturer’s recommendations. Paired-end reads were generated through sequencing on the NovaSeq 6000 platform (Illumina, CA, USA). For quality control, raw reads were trimmed and filtered to remove potential human contamination using Trimmomatic v0.39^60^and Kneaddata (https://github.com/biobakery/kneaddata) respectively, followed by the removal of ribosomal RNA (rRNA) sequences using SortMeRNA v4.3.4^79^. The resulting non-rRNA reads were mapped to the gene sets from corresponding metagenomic samples using HISATA2 v2.2.1^80^ with default parameters. The transcript read counts for each gene were quantified using featureCounts v2.0.1^81^ with default parameters, and subsequently normalized to transcript per million (TPM) per cell using the same method described in the metagenomic analysis above. The gene expression levels, representing the relative number of transcripts per functional gene group, were defined as the ratio between the transcript abundance and the gene abundance in each sample^71^.

### GC-MS analysis of low-molecular-weight aromatic compounds

Gas chromatography-mass spectrometry (GC-MS) was employed to analyze low-molecular-weight (LMW) aromatic compounds in sediment samples. Briefly, 2 g sediment sample were mixed thoroughly with 5 ml of sterile water. The slurries were spiked with ethyl vanillin (a surrogate standard), acidified to pH 1-2 with HCl, and saturated with NaCl prior to three successive 3 mL extractions with ethyl acetate. The extracts were pooled, dehydrated over anhydrous Na_2_SO_4_, dried under nitrogen gas, and derivatized with N,O-bis-(trimethylsilyl)trifluoroacetamide (BSTFA)/pyridine (70°C, 45 min) to generate trimethylsilyl derivativesfor enhanced volatility. 1 µl of the silylated sample was injected into a Trace1310 gas chromatograph coupled to a TSQ8000 mass spectrometer (Thermo Fisher Scientific, USA) equipped with a HP-5MS capillary column (30 m × 0.25 mm i.d.,0.25-μm film thickness). The oven program initiated at 65°C (2 min hold) and ramped at 6 °C min⁻¹ to 300°C (20 min hold). The injector temperature was set at 300℃, and the transfer line and ion source were maintained at 300°C and 290°C, respectively. Helium served as the carrier gas at a flow rate of 1 ml min^-1^. The mass spectrometer operated in the electronimpact (EI) mode at 70 eV, acquiring mass spectra over a range of 50–650 m/z in the full-scan mode. Compound were identified by matching the acquired mass spectra against the NIST library, and quantified relative to the surrogate standard (ethyl vanillin) to correct for compound loss during extraction procedures.

### Microbial bioenergetic modeling

The model implemented in this study divides the microbial community of sediments into four different groups (A1, A2, B1 and B2), which represent archaeal and bacterial lineages with distinct physiological traits and metabolic capabilities, and their differing affinities for relatively labile (OC1) and recalcitrant organic carbon pools (OC2). The specific state variables are listed in Supplementary Method. We assume that the bacterial group B1 and archaeal group A1 utilize labile OC (OC1), while the bacterial group B2 and the archaeal group A2 utilize recalcitrant OC (OC2). *Bathyarchaeia* are represented by archaeal group A2 in this model. We assumed that ∼66% of the organic matter deposited into the seafloor surface of the ECS core is relatively recalcitrant, while the remainder is relatively labile. This assumption is based on the proximities of the sampling sites to the Yangtze River Estuary, which produces a high proportion of the terrigenous input (Supplementary Fig.1) and rapid sedimentation rates^30^.

In this model, a series of ordinary differential equations (ODEs) are used to partition life-history strategies based on bioenergetic assignments among four microbial guilds (B1, B2, A1 and A2). These guilds are characterized by parameters representing maximum growth rate (V_max_), maintenance demand (m_q_), mortality (α), substrate affinity (half-saturation constants, K_V_), growth yield (Y_G_), OC threshold for exogenous maintenance (K_M_) and the ratio of necromass to OC (See details in Supplementary Methods). This parameterization reflects the physiological trade-off between rapid growth and long-term persistence under energy limitation in the deep subsurface. The rate of substrate uptake (𝑈*_i_*) for each microbial guild 𝑖 ∈ {𝐵_1_, 𝐵_1_, 𝐴_1_, 𝐴_2_} is governed by standard Michaelis-Menten kinetics (𝐾*_V_*) depending on the specific OC concentration (𝐶*_j_*):

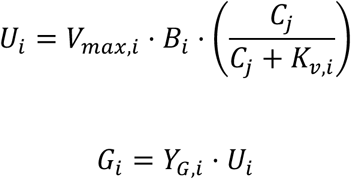

where 𝑉*_max,i_* is the maximum uptake rate, is the biomass, and 𝐾*_v,i_* is the half-saturation constant for uptake. The actual synthesized biomass for each microbial guild (𝐺*_i_*) 𝑖 ∈ {𝐵_1_, 𝐵_2_, 𝐴_1_, 𝐴_2_} is scaled by their true growth yields (𝑌*_G,i_*).

To mechanistically resolve microbial survival under energy limitation in the deep biosphere, we introduced a dynamic switch. The model conceptualizes that basal maintenance energy shifts from exogenous OC assimilation to endogenous cellular catabolism as external energy wanes. This physiological state transition is regulated by a Fermi-Dirac sigmoidal function (𝜃*_M,i_*):

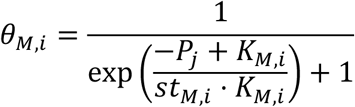

where 𝐾*_M,i_* is the threshold of OC concentration regulating the provenance of maintenance power, and 𝑠𝑡*_M,i_* determines the steepness of the state change. The exogenous maintenance (𝑀*_Ex,i_*) and endogenous maintenance (𝑀*_En,i_*) rates are subsequently defined as:

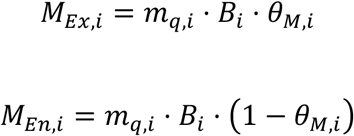

The overall model conceptualization, formulations, parameterization and other details are presented in the Supplementary Methods. The simulation spans a 1,000-year burial timescale based on the documented sedimentation rate of 0.61 cm yr⁻¹ and a dry bulk density of 1.1 g cm⁻³ in ECS shelf^30^. Microbial biomass was converted to carbon units using a cell-specific content of 24 fg C cell^-13^. Initial conditions and parameters were constrained by empirical observational data from the core A2 in this study are provided in the Supplementary Methods. The model was implemented in the R programming language and numerically integrated using the lsoda solver from the deSolve package^82^.

### Global budget and flux of OC degradation

We calculated the global transfer efficiency of OC over the past 100 ka ^5^. The time-integrated OC flux into global shelf and marginal regions was used to estimate the total accumulated organic carbon during the past 1,000 years. The proportions of recalcitrant OC and the weighted coefficients of OC degradation rates mediated by A2 (*Bathyarchaeia*) are derived from the modeling results, thus enabling us to quantify the contribution of Bathyarchaeia to OC degradation in global shelf sediments over millennial timescales of sediment burial.

## Data availability

16S rRNA gene amplicon, metagenomic and metatranscriptomic raw data are available at NCBI under bioproject PRJNA1434522, the 38 *Bathyarchaeia* MAGs analyzed are available at eLMSG (https://biosino.org/elmsg/index) under accession numbers LMSG_G000011401.1 to LMSG_G000011440.1.

## Code availability

Data analysis pipelines used in this work can be obtained from https://github.com/houjialin/ECS-Bathy-Carbon-model.git

## Acknowledgements

We thank Professor Bo Barker Jørgensen for constructive comments that significantly improved the manuscript. This work is supported by Natural Science Foundation of China (Grants 42230401,42406089), 2030 Project of Shanghai Jiao Tong University (Grant WH510244001) and the project Ocean Negative Carbon Emission (ONCE); JLH was supported by the Postdoctoral Fellowship Program (Grade B) of CPSF (GZB2024030); JAB was supported by the Agence Nationale de la Recherche (ANR23-CPJ1-0172-01) and the European Research Council (ERC) under the European Union’s Horizon Europe Research and Innovation programme (Grant agreement No. 101115755, acronym SIESTA).

## Contributions

F.P.W conceptualized, supervised and provided major funding and resources to this work. L.W.L led the sample collection during the ECS cruise supported by F.P.W and Z.J.W and generated the microbial, geochemical and multi-omics data with assistance from W.K.S, L.Y.L and H.N.H. L.W.L, L.Y.L and J.L.H performed the qPCR and 16S rRNA genes analysis. J.L.H performed the meta-omics data analysis and modeling with assistance from J.A.B and L.Z. L.D. L.H.D and O.S edited and reviewed the final manuscript. J.L.H and L.W.L wrote the original draft manuscript and prepared the revision with input from all authors.

## Extended Data figures

**Extended Data Fig.1.**
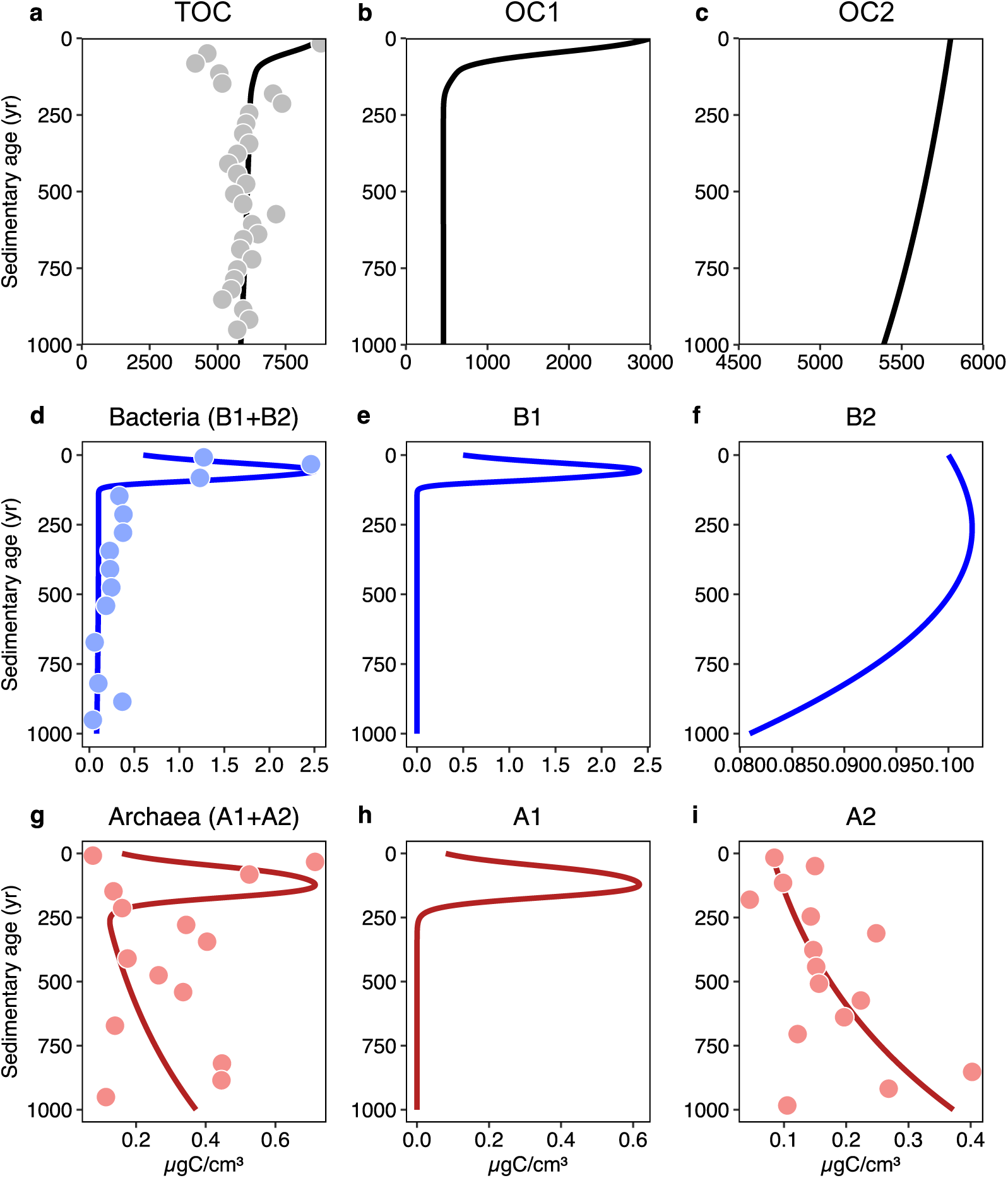
Comparison of observed and simulated organic carbon pools and microbial biomass in the sediment core A2. Model simulations are shown as lines, and empirical observations are represented by dots. **a**, total organic carbon (OC). **b**, simulated labile OC1 concentration. **c**, simulated recalcitrant OC2 concentration. **d**, total biomass of bacteria. **e**, simulated biomass of bacterial guild B1. **f**, simulated biomass of bacterial guild B2. **g**, total biomass of archaea. **h**, simulated biomass of archaeal guild A1. **i**, the biomass of archaeal guild A2 represented by *Bathyarchaeia*.

**Extended Data Fig.2.**
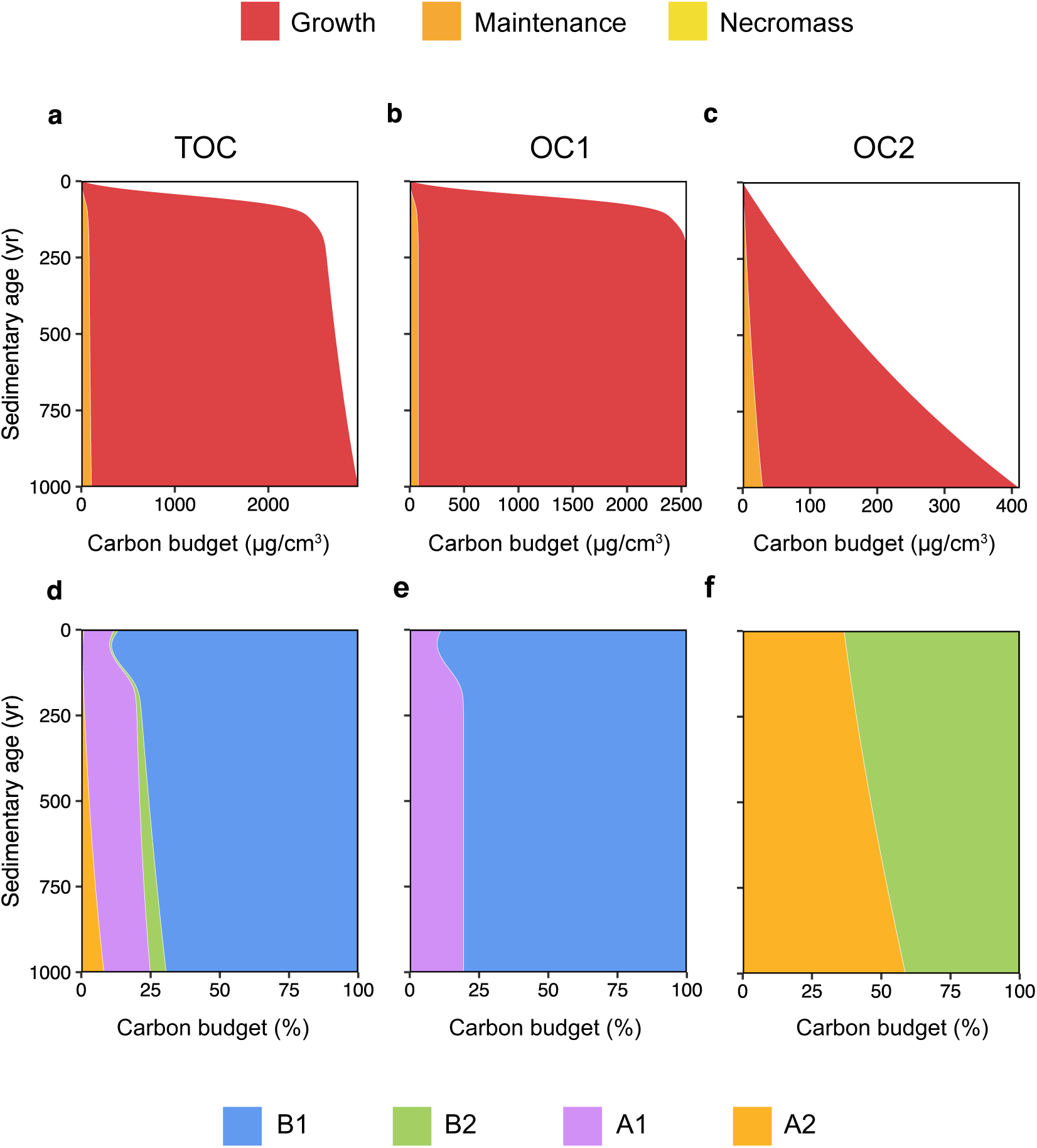
Simulated microbial contribution to OC budgets over a 1,000-year timescale. Simulated carbon budgets partitioning into microbial growth, exogenous maintenance, and necromass generated from mortality for **a**, total OC (TOC), **b**, labile OC1, **c**, recalcitrant OC2. **d-f**, stacked charts of relative contributions (%) from each microbial guild to the carbon budget of **d**, TOC, **e**, labile OC1, and **f**, recalcitrant OC2. Guild colors are blue (B1), green (B2), purple (A1) and orange (A2). Panel distributions reflect distinct substrate preferences, where labile OC2 is utilized by B1 and A1 guilds and recalcitrant OC2 by B2 and A2 guilds.

**Extended Data Fig.3.**
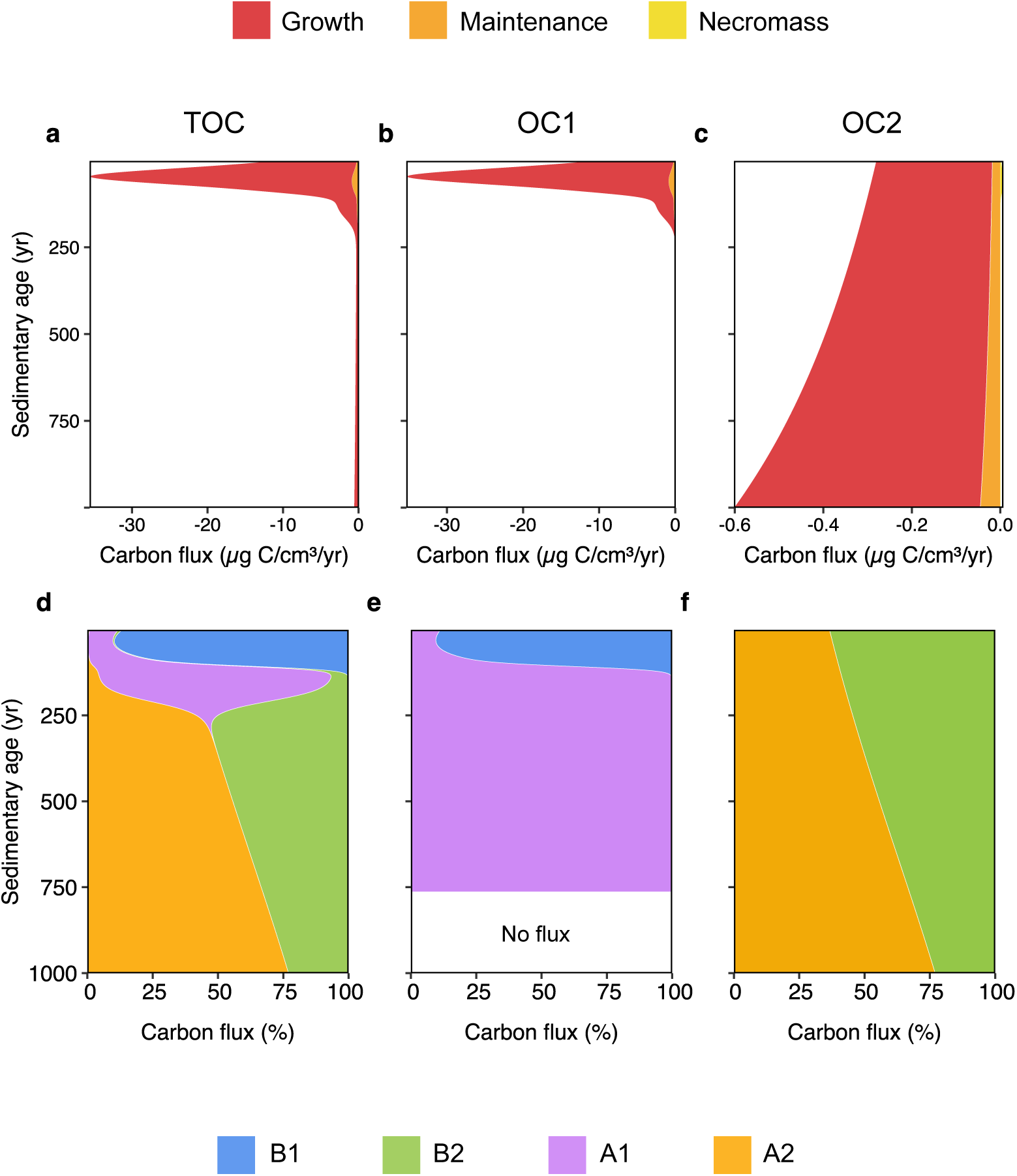
Simulated microbial carbon fluxes to OC pools over a 1,000-year timescale. **a–c**, Simulated carbon fluxes partitioning into microbial growth (red), exogenous maintenance demand (orange), and necromass generated from mortality (bright yellow) for **a**, TOC, **b**, labile OC1, and **c**, recalcitrant OC2. **d–f**, Stacked charts presenting relative contributions (%) of four microbial guilds to the carbon fluxes for **d**, TOC, **e**, labile OC1, and **f**, recalcitrant OC2. Guild colors are blue (B1), green (B2), purple (A1) and orange (A2). Panel distributions reflect distinct substrate preferences defined in the model, where labile OC1 flux is utilized exclusively by B1 and A1 guilds and recalcitrant OC2 flux by B2 and A2 guilds. Label “No flux” in **e** indicates complete not consumption of labile OC1 and cessation of associated microbial activity.

**Extended Data Fig.4.**
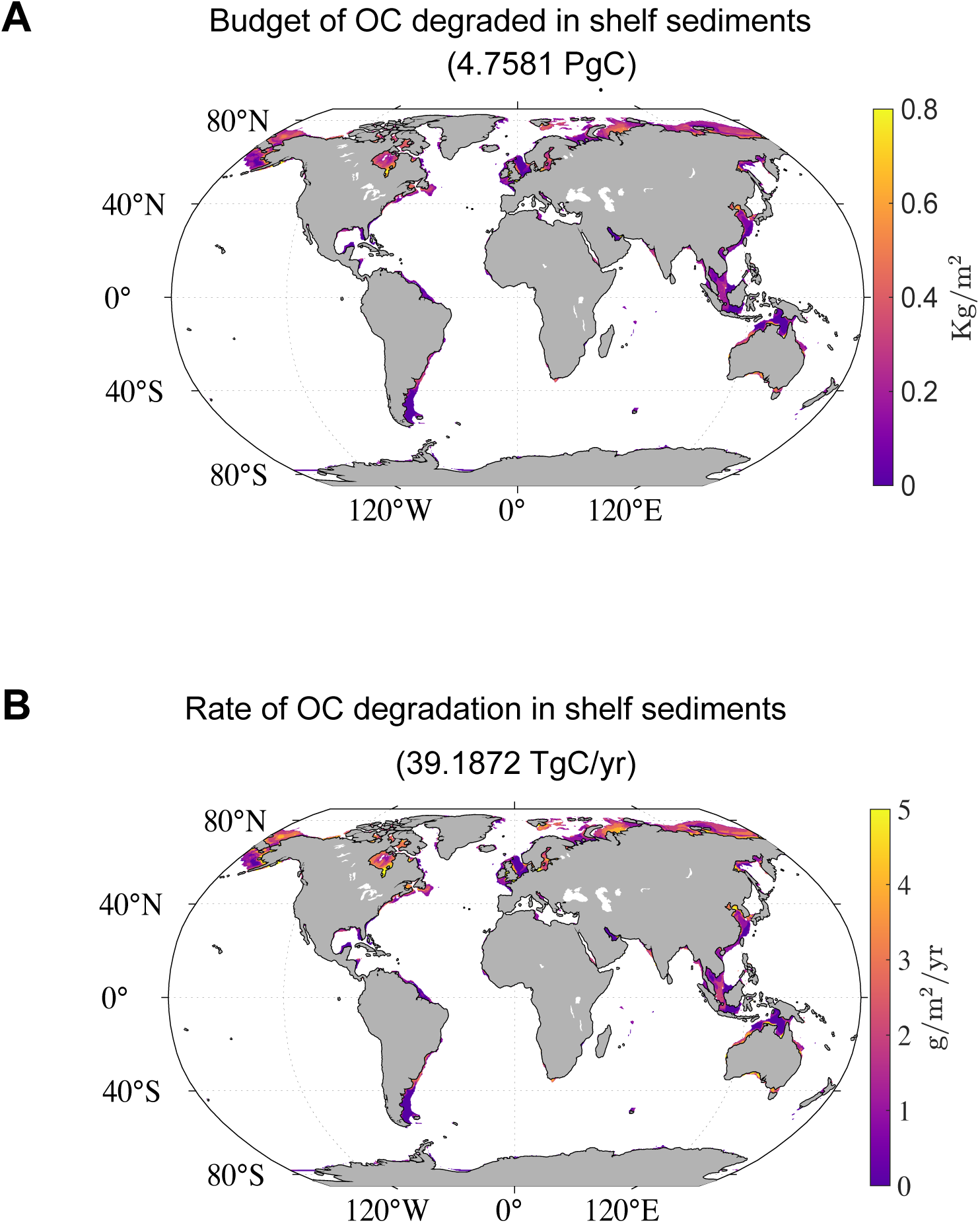
Global budget and rate of particulate organic carbon degradation mediated by Bathyarchaeia in global shelf sediments over 1,000 years of burial. **a**, Global distribution of the total budget (4.7581 PgC) of POC degraded in shelf sediments over millennial timescales. **b**, Global distribution of the rate of POC degradation (39.1872 Tg C/yr) in shelf sediments after 1000 years burial.

**Extended Data Fig.5.**
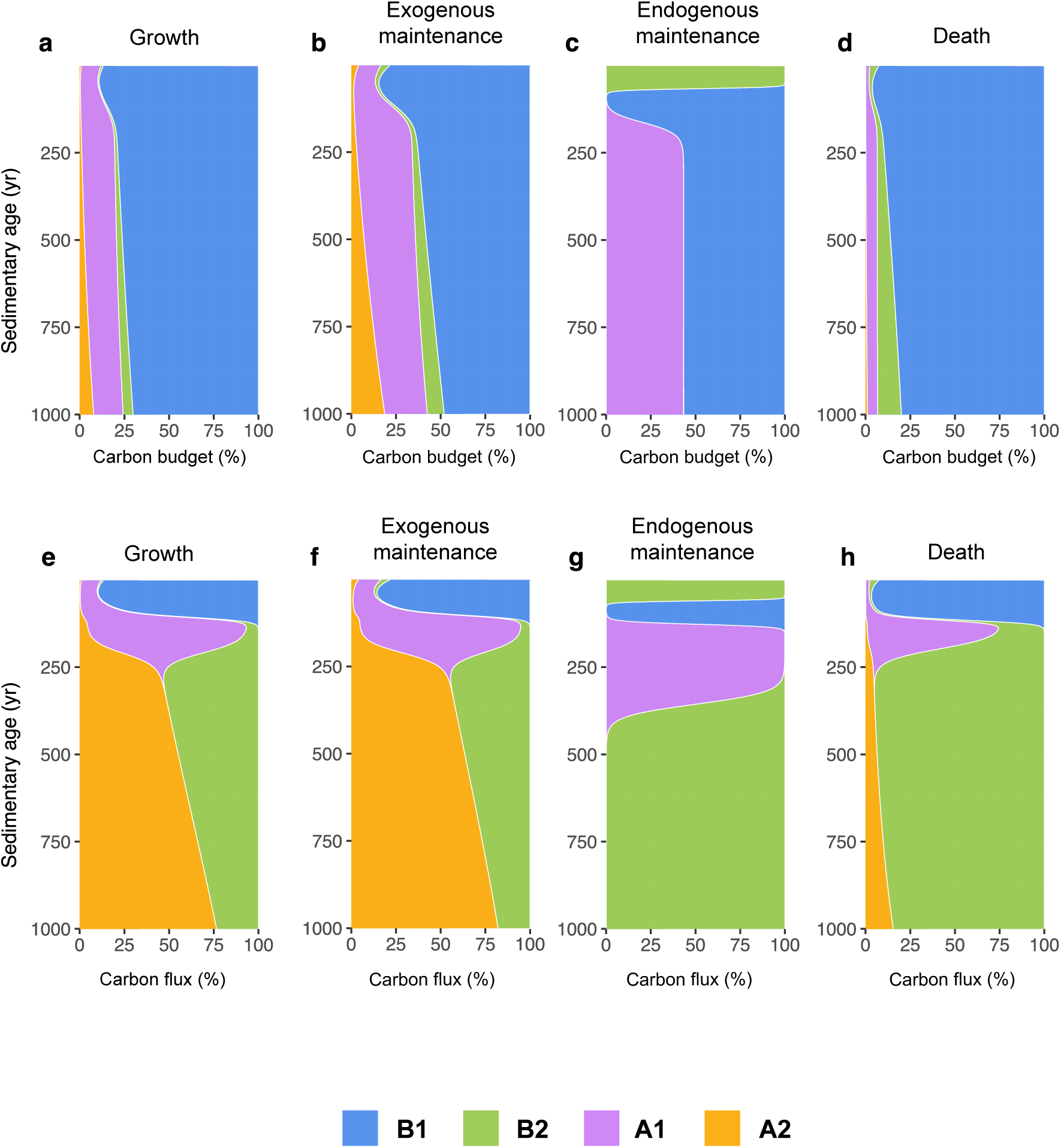
Dynamic changes of OC budget and flux among different microbial guilds through various physiological metabolic activities over a millennial timescale. **a-d**, Panels show the relative contribution (%) of carbon budget pools for different physiological activities among the four microbial guilds. Carbon budgets are assigned to: **a**, Microbial growth. **b**, Exogenous maintenance demand (maintenance energy derived from external OC). **c**, Endogenous maintenance demand (maintenance energy derived from consuming internal cellular biomass). **d**, Necromass generated via mortality. **e–h,** Panels show the relative contribution (%) of carbon flux rates for: **e**, Microbial growth rate. **f**, Exogenous maintenance flux. **g**, Endogenous maintenance flux. **h**, Mortality

## Supplementary information

Supplementary Methods. The bioenergetic modeling conceptualization, formulations, parameterization and implementation

Supplementary Table 1. The sampling information of six sediment cores from East China Sea continental shelf.

Supplementary Table 2. The qPCR results of 16S rRNA genes for Bacteria and Archaea in the six sediment cores.

Supplementary Table 3. The analysis result of archaeal amplicon sequencing in the six sediment cores.

Supplementary Table 4. The global amplicon sequencing dataset and sampling information used in the Fig.2B.

Supplementary Table 5. Calculated doubling time of different microbial population in the sediment of core A2.

Supplementary Table 6. The functional annotation of key genes in this study.

Supplementary Table 7. Genomic information of 38 high- and median-quality MAGs recovered from five metagenomes of core A2.

Supplementary Fig.1 The proportion of terrestrial organic carbon that calculated based on the carbon isotopic composition of bulk organic matter (δ^13^C_org_) contribute to the total organic carbon in the sediment cores **a**, A2, **b**, A3, **c**, A5, **d**, B1, **e**, B2 and **f**, B3 from East China Sea continental shelf. The sources of TOC contents were constrained using a two end-member mixing model following the Equation: 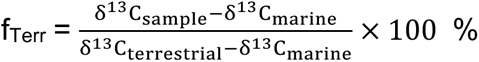, f_Terr_ represents the contribution of terrestrial source to TOC, δ^13^C_sample_ represents the measured values of sediment samples, while δ^13^C_marine_ and δ^13^C _terrestrial_ represent the end member values of marine- and terrestrial-derived TOC, respectively. For δ^13^C_marine_, a value of -20‰ was adopted^83^, while for δ^13^C_terrestrial_, a value of -26‰ was used^84^.

Supplementary Fig.2 Geochemical profiles (Sulfate, methane, DIC, δ^13^C-DIC, TOC, δ^13^C-TOC, Ca^2+^, Mg^2+^ and Cl^-^) of porewater in the sediment cores of **a**, A2, **b**, A3, **c**, A5, **d**, B1, **e**, B2 and **f**, B3 from East China Sea continental shelf.

Supplementary Fig.3 The gene abundance, transcript abundance, expression levels and taxonomic composition of key genes from the metagenomes of five depth in A2 core.

Supplementary Fig.4 The concentration of aromatic compounds measured in the sediment core A2. **a**, Benzene-1,3-bis (1,1-dimethylethyl), **b**, 2,4-Di-tert-butylphenol, **c**, Cholesterol, **d**, 2-Hydroxylamino-4,6-dinitrotoluene, **e**, β-Sitosterol, **f**, Stigmastanol, **g**, 4α,23,24-Trimethyl-5α-cholest-22-en-3β-ol.

Supplementary Fig.5 The gene and transcript abundance of the genes involved in microbial antioxidation mechanism in the sediment core A2, including **a**, thioredoxin-dependent peroxiredoxin [EC:1.11.1.24] (BCP, K03564), **b**, superoxide dismutase, Fe-Mn family [EC:1.15.1.1] (SOD2, K04564), **c**, superoxide reductase [EC:1.15.1.2] (dfx, K05919).

