## Supplementary figures and images for "Physiological trade-offs drive the archaeal dominance and carbon turnover in deep subsurface"

### Supplementary Figure 1

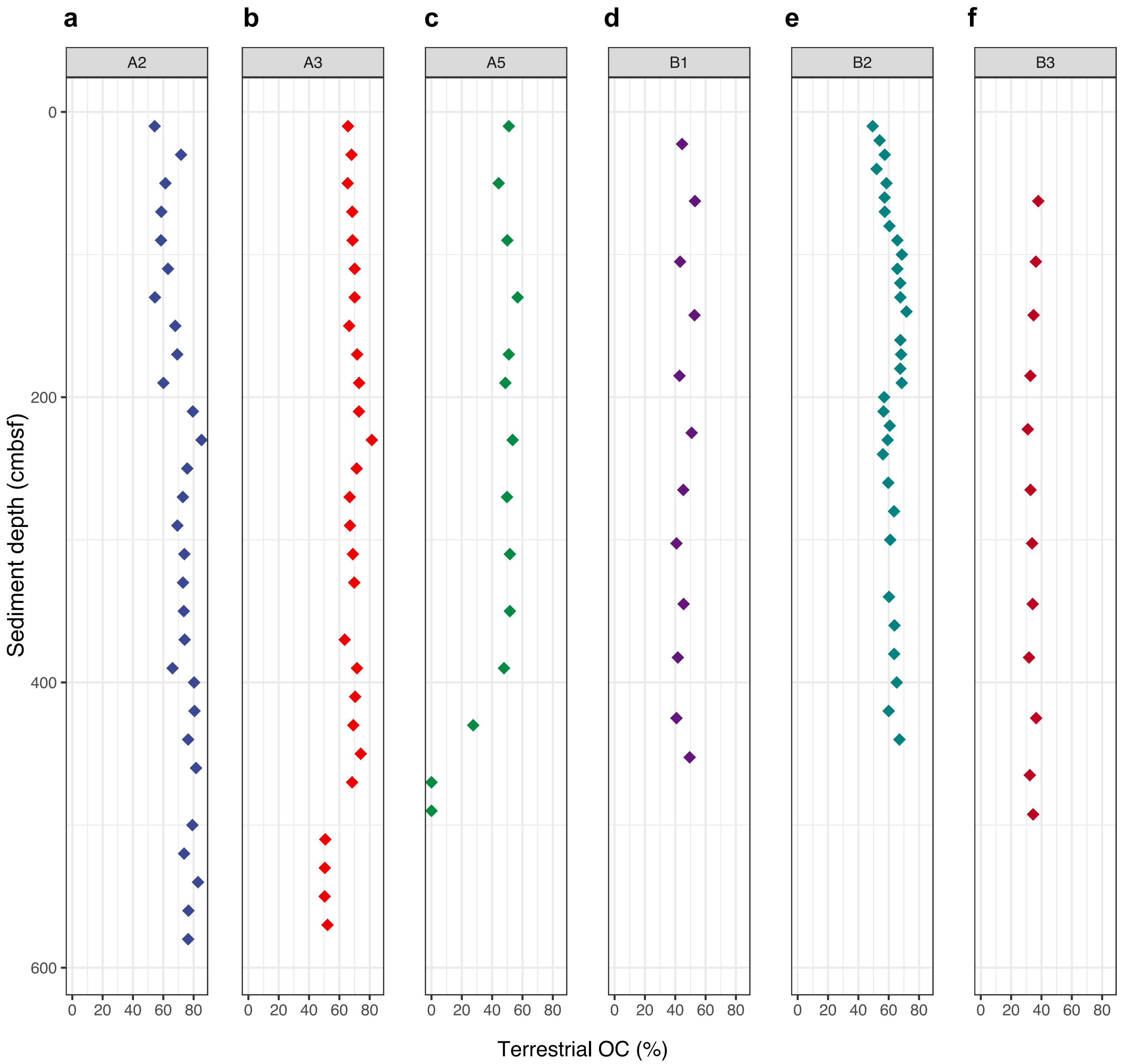

### Supplementary Figure 2

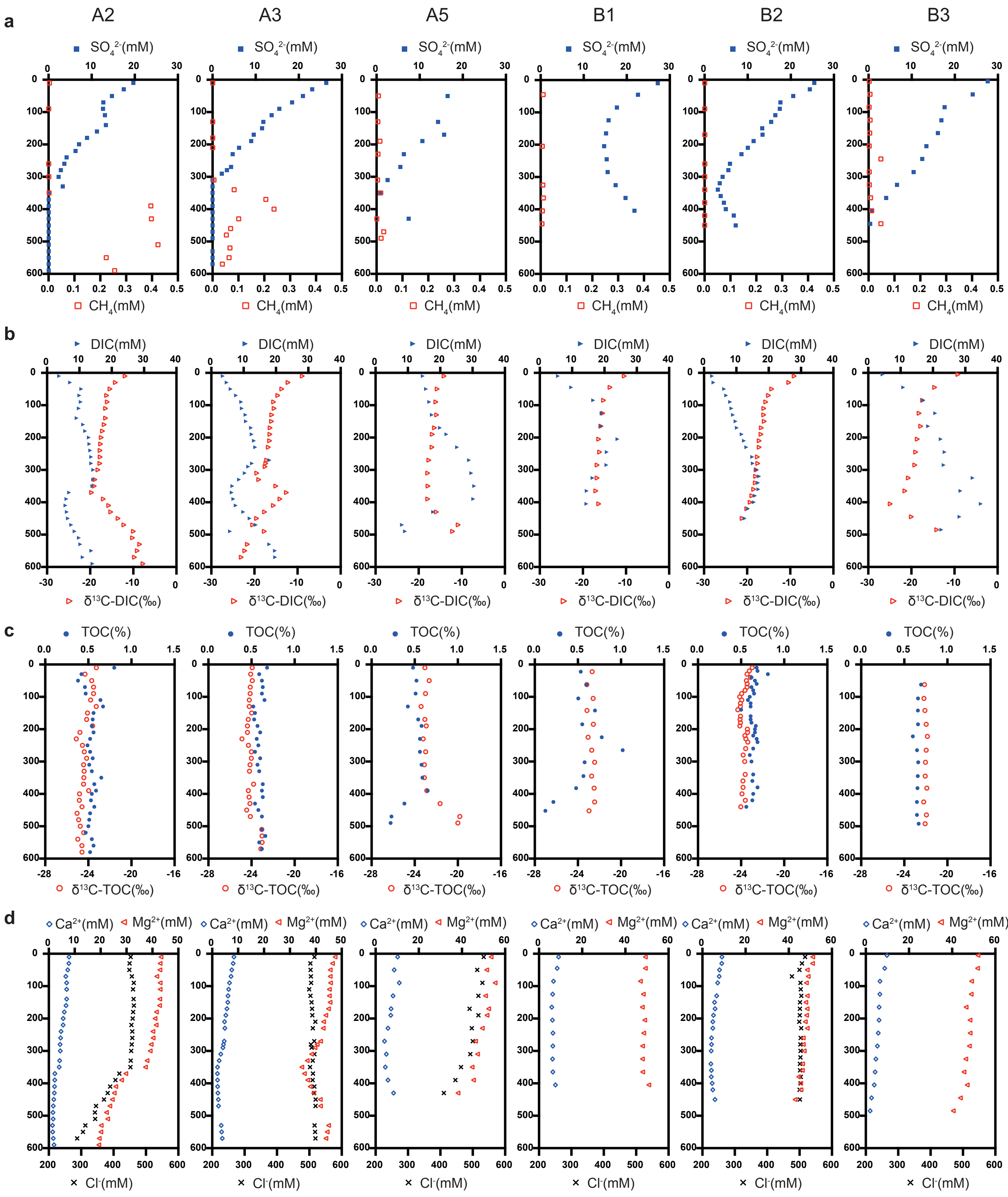

### Supplementary Figure 3

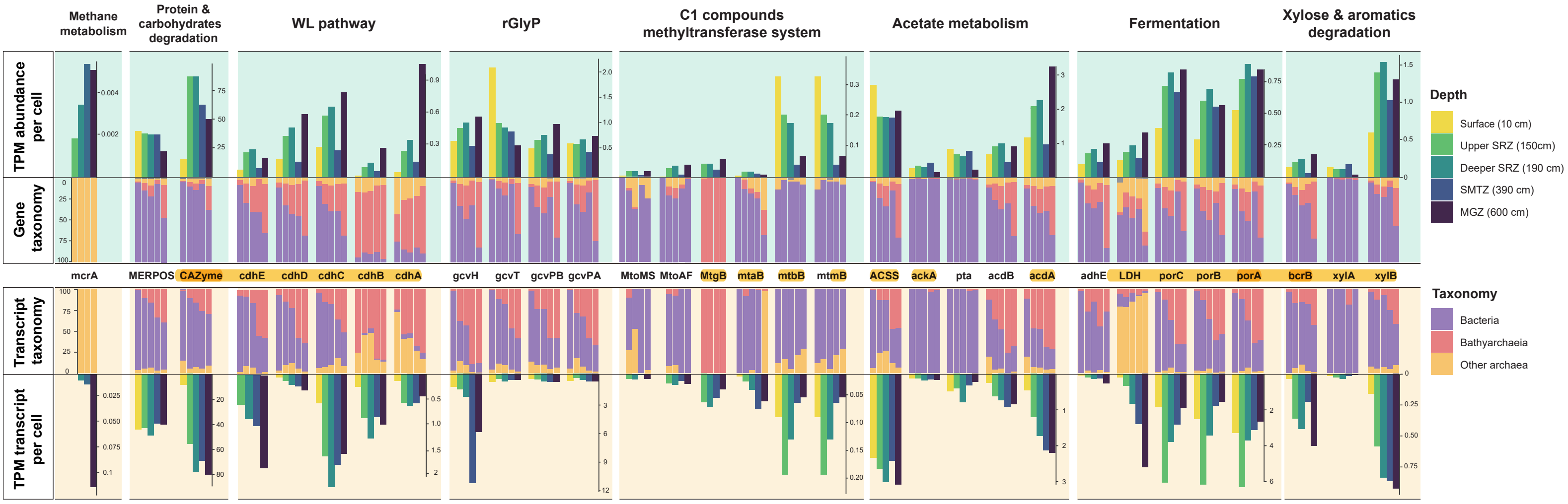

### Supplementary Figure 4

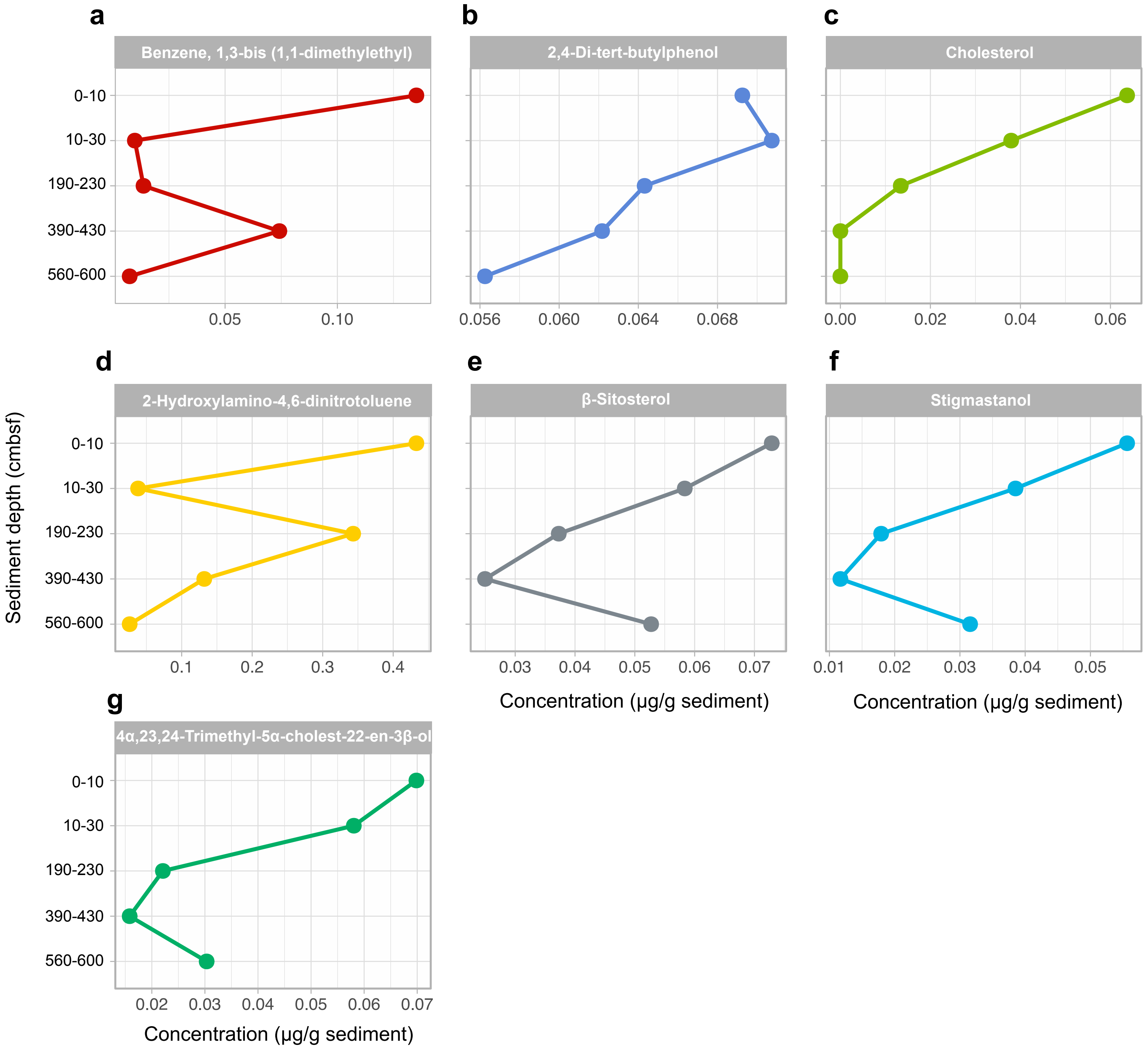

### Supplementary Figure 5

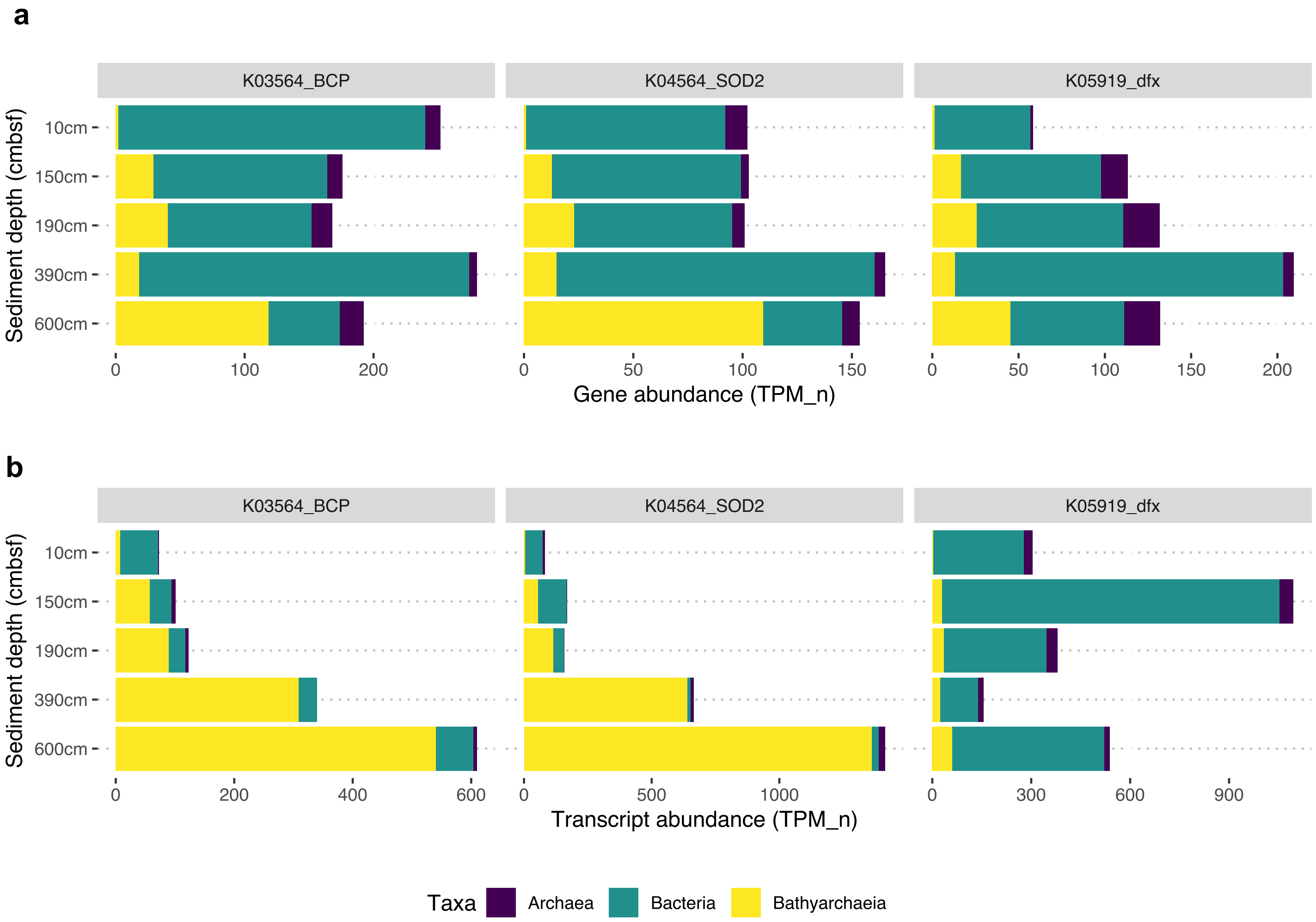
