## Supplementary Methods for "Physiological trade-offs drive the archaeal dominance and carbon turnover in deep subsurface"

### 1. Geochemical characterization of the sediment cores

Upon retrieval, porewater of sediment cores was extracted using Rhizon samplers (0.2-μm pore size; Rhizosphere). Subsequently, sediments for microbiological analysis were stored at -80°C until DNA extraction. For A2, A3, A5 and B2, sulfate (SO42-), Cl-, Ca2+, Mg2+ methane (CH4), dissolved inorganic carbon (DIC), 13C-DIC, were measured as described previously(Liu et al., 2020). Briefly, SO42- and Cl-concentration were measured by ion chromatography (ICS-1500, DIONEX) equipped with a 4 mm AS-9HC column. CH4 was measured by a gas chromatograph with flame ionization detector (GC-FID, Agilent). DIC concentrations and δ13C-DIC values were measured using a Gas Bench II isotope-ratio mass spectrometer (Gas Bench II-IRMS, Thermo Fisher Scientific). Ca2+ and Mg2+ were measured using inductively coupled plasma atomic emission spectroscopy (ICP-AES; IRIS Advantage, Thermo Jarrell Ash). For B1 and B3(Israel), dissolved sulfate using a Dionex DX500 high-performance liquid chromatograph (HPLC) with an error of 3%. Headspace methane concentrations were measured on a Thermo Scientific gas chromatograph (GC) equipped with a flame ionization detector (FID) at a precision of 2 μmol·L-1. Sulfate concentrations were analyzed by inductively coupled plasma-atomic emission spectroscopy (ICP-OES-720-ES, VRIAN) with a precision of 2% . Total organic carbon and δ13C-TOC were measured by elemental analyser (Vario EL III, Elementar) coupled with IRMS (Isoprime, Elementar) at the instrumental analysis center, Shanghai Jiao Tong University.

### 2. Model Conceptualization

To mechanistically resolve the survive strategy behind the subsurface microbial succession and quantitatively evaluate their roles in the degradation of sedimentary organic carbon (OC) over geological timescales, we developed a bioenergetic model that integrated microbial ecophysiology and long-term kinetics of OC mineralization.

#### 2.1 The 2-G carbon mineralization model

This model partitions the total organic carbon (TOC) into two discrete pools based on their reactivity: a labile pool ($OC_1$) and a recalcitrant pool ($OC_2$).

- $\boldsymbol{OC_1}$ **(Labile OC Pool):** Represents relatively fresh and high-bioavailable organic matter that shortly deposited into the seafloor from the overlying seawater.

- $\boldsymbol{OC_2}$ **(Recalcitrant OC Pool):** Comprises complex, refractory compounds—such as terrigenous aromatic lignin—that dictate the long-term carbon reactivity and low burial efficiency over millennial timescales.

##### 2.2 Ecophysiological guilds and life-history strategy

The microbial community is classified into four distinct ecophysiological guilds, distinguished by their physiological traits and substrate preferences to the two OC pools:

- **Bacteria** $B_1$: Typical copiotrophic bacteria specializing in their high maximum grwoth rates ($V_{max}$), growth yield ($Y_G$) and high mortality ($\alpha$), these bacteria prioritize rapid biomass turnover using the labile $OC_1$ but incur substantial maintenance costs ($m_q$)

- **Bacteria** $B_2$: Generalist bacteria primarily targeting recalcitrant $OC_2$ with reduced maximum growth rate, manitenance costs compared to the typical copiotrophic $B_1$.

- **Archaea** $A_1$: Utilizing labile $OC_1$, this guild adopts a more Moderately growth-maintenance strategy, allowing for persistence slightly deeper than $B_1$.

- **Archaea** $A_2$: Extremely oligotrophic archaea, specifically represented by Bathyarchaeia in this model, adapted to the deep biosphere, characterized by ultra-low mortality, slow maximum growth rate, low maintenance energy demands, crucially, $A_2$ possesses a superior substrate affinity to the recalcitrant $OC_2$ (low half-saturation constant $K_V$), enabling it to outcompete $B_2$ in the energy-limited deep subsurface environments.

#### 3. Model Formulation

A system of coupled ordinary differential equations (ODEs) describes the transfers and transformations of carbon among the four microbial guilds and two OC pools, driven by microbial substrate uptake, growth, maintenance, mortality, and necromass recycling.

##### 3.1 Substrate Uptake and Microbial Growth

We assume that OC degradation is strictly coupled to the specific microbial substrate consumption. The rate of substrate uptake ($U_i$) for each microbial guild $i \in \{B_1, B_2, A_1, A_2\}$ is governed by standard Michaelis-Menten kinetics ($K_V$) depending on the specific OC concentration ($C_j$):

$$U_i = V_{max,i} \cdot B_i \cdot \left(\frac{C_j}{C_j + K_{v,i}}\right)$$

where $V_{max,i}$ is the maximum uptake rate, $B_i$ is the biomass, and $K_{v,i}$ is the half-saturation constant for uptake. The actual synthesized biomass for each microbial guild $(G_i)$ $i \in \{B_1, B_2, A_1, A_2\}$ is scaled by their true growth yields $(Y_{G,i})$:

$$G_i = Y_{G,i} \cdot U_i$$

##### 3.2 Maintenance Energy and Physiological State

To survive in the deep subsurface, microorganisms must meet basal maintenance power demands $(m_{q,i})$ required for their cellular integrity and biomolecular repair under energy-limited and other extreme conditions. The model permits flexibility in the provenance of this maintenance energy: exogenous (derived directly from OC) or endogenous (derived from the catabolism of their own cellular biomass). Following the theoretical framework of Stolpovsky et al. (2011) and Bradley et al. (2019), the transition between exogenous and endogenous maintenance is controlled by a sigmoidal function $(\theta_{M,i})$ based on Fermi-Dirac statistics:

$$\theta_{M,i} = \frac{1}{\exp\left(\frac{-P_j + K_{M,i}}{st_{M,i} \cdot K_{M,i}}\right) + 1}$$

where $K_{M,i}$ is the threshold of OC concentration regulating the provenance of maintenance power, and $st_{M,i}$ determines the steepness of the state change. The exogenous maintenance $(M_{Ex,i})$ and endogenous maintenance $(M_{En,i})$ rates are subsequently defined as:

$$M_{Ex,i} = m_{q,i} \cdot B_i \cdot \theta_{M,i}$$

$$M_{En,i} = m_{q,i} \cdot B_i \cdot \left(1 - \theta_{M,i}\right)$$

Under non energy-limited conditions with high OC content $(P_j > K_{M,i})$, $\theta_{M,i}$ approaches 1, and microbial maintenance is supported exogenously by OC. While OC is depleted below $K_{M,i}$ with increasing burial age, this model simulates a transition where maintenance requirements are increasingly met by consuming internal biomass (endogenous) rather than external OC (exogenous).

##### 3.3 Mortality and Necromass Recycling

Microbial mortality $(D_i)$ follows first-order kinetics defined by a guild-specific mortality rate $(\alpha_i)$:

$$D_i = \alpha_i \cdot B_i$$

A fraction ($f_{recyc}$) of the total dead biomass ($\Sigma D_i$) is recycled back into the system as necromass. This recycled organic matter is partitioned into the labile $OC_1$ and refractory $OC_2$ pools based on a fractionation coefficient ($f_{POC1}$):

$$D_{to\_POC1} = f_{recyc} \cdot f_{POC1} \cdot \sum D_i$$

$$D_{to\_POC2} = f_{recyc} \cdot (1 - f_{POC1}) \cdot \sum D_i$$

### 4. Carbon mass balance equations (ODE system)

Integrating the aforementioned bioenergetic fluxes, the governing equations for the four microbial biomass ($B_i$) and two OC pools ($OC_1, OC_2$) are formulated as follows:

For biomass pools of four microbial guilds:

$$\frac{dB_1}{dt} = G_{B1} - M_{En,B1} - D_{B1}$$

$$\frac{dB_2}{dt} = G_{B2} - M_{En,B2} - D_{B2}$$

$$\frac{dA_1}{dt} = G_{A1} - M_{En,A1} - D_{A1}$$

$$\frac{dA_2}{dt} = G_{A2} - M_{En,A2} - D_{A2}$$

For labile and recalcitrant CO pools:

$$\frac{dPOC_1}{dt} = D_{to\_POC1} - (U_{B1} + U_{A1}) - \left(M_{Ex,B1} + M_{Ex,A1}\right)$$

$$\frac{dPOC_2}{dt} = D_{to\_POC2} - (U_{B2} + U_{A2}) - \left(M_{Ex,B2} + M_{Ex,A2}\right)$$

### 5. Parameterization and Implementation

The model simulates a 1,000-year timescale, converting burial depth to age using a sedimentation rate $\omega = 0.61$ cm yr$^{-1}$ and a bulk dry density $\rho = 1.1$ g cm$^{-3}$ based on the reported data from near sites (Yang et al., 2023). Cell density is converted to biomass using an average cell dry weight 24 fg C/cell (Bar-on et al., 2018). Initial conditions and parameters were constrained by empirical observational data from the A2 site in this study and recent bioenergetic estimations (Bradley et al, 2019) (Table S1 and S2). The model was implemented in the R programming language and numerically integrated using the lsoda solver from the deSolve package.

#### Table S1 Model Parameters

| Parameter | Description | Unit | $B_1$ | $B_2$ | $A_1$ | $A_2$ | Reference |
|---|---|---|---|---|---|---|---|
| $V_{max}$ | Maximum growth rate | $yr^{-1}$ | 60 | 4 | 50 | 2 | Bradley et al., 2019 |
| $K_v$ | Half-saturation constant | $\mu g\ C\ cm^{-3}$ | 6300 | 8000 | 6800 | 4000 | This study |
| $Y_G$ | True growth yield | unitless | 0.006 | 0.012 | 0.003 | 0.002 | This study |
| $\alpha$ | Mortality rate | $yr^{-1}$ | $7 \times 10^{-2}$ | $2 \times 10^{-2}$ | $1 \times 10^{-2}$ | $8 \times 10^{-4}$ | Bradley et al., 2019 |
| $m_q$ | maintenance demand | $yr^{-1}$ | 0.3 | 0.1 | 0.3 | 0.1 | Bradley et al., 2019 |
| $K_M$ | OC threshold of maintenance state change | $\mu g\ C\ cm^{-3}$ | 600 | 3000 | 400 | 2000 \| | This study |
| $st_M$ | Steepness of state-change dependency | unitless | 0.1 | 0.1 | 0.1 | 0.1 | Bradley et al., 2019 |

#### Table S2 Initial conditions

| Initial conditions | Description | Unit | Value | Observation |
|---|---|---|---|---|
| Total OC | The total OC content in the 0-2 cm of sediments in A2 site | $\mu g\ C\ cm^{-3}$ | 8800 | 0.8% wet weight<br>(Convert to $\mu g\ C\ cm^{-3}$ based on the sediment density) |
| Labile $OC_1$ | The labile OC buried into the seafloor | $\mu g\ C\ cm^{-3}$ | 5800 | Nearly 66% of Total OC |
| Recalculate $OC_2$ | Recalculate OC buried into the seafloor | $\mu g\ C\ cm^{-3}$ | 3000 | Nearly 34% of Total OC |

| | | | | |
|---|---|---|---|---|
| Total biomass | The initial total biomass in the seafloor | $\mu$g C cm$^{-3}$ | 1.35 | Total cell density estimated based on qPCR of 16S rRNA gene |
| B1 | The initial biomass of B1 in the seafloor | $\mu$g C cm$^{-3}$ | 0.5 | - |
| B2 | The initial biomass of B2 in the seafloor | $\mu$g C cm$^{-3}$ | 0.1 | - |
| A1 | The initial biomass of A1 in the seafloor | $\mu$g C cm$^{-3}$ | 0.08 | - |
| A2 | The initial biomass of A2 in the seafloor | $\mu$g C cm$^{-3}$ | 0.08 | 0.08 (Estimated Bathyarchaeia biomass at 10 cmbsf) |
| $f_{recyc}$ | The fraction of dead cell converted to necromass | unitless | 0.4 | - |
| $f_{OC1}$ | The fraction of necromass converted to labile OC1 | unitless | 0.9 | - |
